# scMINER: a mutual information-based framework for identifying hidden drivers from single-cell omics data

**DOI:** 10.1101/2023.01.26.523391

**Authors:** Liang Ding, Hao Shi, Chenxi Qian, Chad Burdyshaw, Joao Pedro Veloso, Alireza Khatamian, Qingfei Pan, Yogesh Dhungana, Zhen Xie, Isabel Risch, Xu Yang, Xin Huang, Lei Yan, Michael Rusch, Michael Brewer, Koon-Kiu Yan, Hongbo Chi, Jiyang Yu

## Abstract

The sparse nature of single-cell omics data makes it challenging to dissect the wiring and rewiring of the transcriptional and signaling drivers that regulate cellular states. Many of the drivers, referred to as “hidden drivers”, are difficult to identify via conventional expression analysis due to low expression and inconsistency between RNA and protein activity caused by post-translational and other modifications. To address this issue, we developed scMINER, a mutual information (MI)-based computational framework for unsupervised clustering analysis and cell-type specific inference of intracellular networks, hidden drivers and network rewiring from single-cell RNA-seq data. We designed scMINER to capture nonlinear cell-cell and gene-gene relationships and infer driver activities. Systematic benchmarking showed that scMINER outperforms popular single-cell clustering algorithms, especially in distinguishing similar cell types. With respect to network inference, scMINER does not rely on the binding motifs which are available for a limited set of transcription factors, therefore scMINER can provide quantitative activity assessment for more than 6,000 transcription and signaling drivers from a scRNA-seq experiment. As demonstrations, we used scMINER to expose hidden transcription and signaling drivers and dissect their regulon rewiring in immune cell heterogeneity, lineage differentiation, and tissue specification. Overall, activity-based scMINER is a widely applicable, highly accurate, reproducible and scalable method for inferring cellular transcriptional and signaling networks in each cell state from scRNA-seq data. The scMINER software is publicly accessible via: https://github.com/jyyulab/scMINER.

## Main

Cell fate determination and specification are governed by the wiring and rewiring of characteristic proteins including transcription factors (TFs) and upstream signaling factors (SIGs). Systematic identification of these cell-type specific drivers is crucial to understanding the cellular plasticity and dynamics, and providing therapeutic targets for diseases^1^. Nevertheless, many drivers, especially SIG drivers, can undergo activity alteration at the posttranslational level without drastic changes in their gene expression level, making them “hidden drivers” and difficult to capture by differential expression analysis. Network-based systems biology algorithms such as NetBID^2^ have been developed to uncover hidden drivers from bulk omics data. However, computational algorithms to infer cell-type specific hidden drivers and network rewiring from single-cell omics data are lacking.

The use of single-cell RNA-sequencing (scRNA-seq) methods have revolutionized our ability to identify cell states with unprecedented resolution. The scRNA-seq data also provide opportunities to dissect the wiring and rewiring of cell states and identify the underlying TF and SIG drivers. However, inherent stochasticity and sparsity arising from variations and fluctuations among genetically identical cells, as well as low signal-to-noise resulting from the heterogeneity within populations of genetically similar cells present unique challenges in network and driver activity inference^3–5^. A number of recent methods have been proposed to reconstruct TF regulatory networks from scRNA-seq data. For example, one of the most commonly used methods, SCENIC^6^, uses TF binding motif databases and co-expression analysis to reconstruct TF-target networks and infer TF activity that can be used for clustering analysis. Although SCENIC is broadly applicable for analysis of scRNA-seq data and inferring TF regulatory networks (TRNs) that define a cell state, it is restricted to the analysis of TF activity alone. Also, the TF cis-regulatory motif databases used for SCENIC are context-independent and incomplete, thus limiting the performance of this methodology. Furthermore, a recent benchmarking analysis of the state-of-the-art methods for TRN inference from scRNA-seq data revealed that all the algorithms analyzed have important limitations^7^, leaving an ongoing demand for robust tools. Additionally, there are currently no algorithms that can infer cell-type specific signaling networks from single-cell transcriptomics data.

Another limitation is to accurately estimate cell-cell similarity and gene-gene dependency, which is critical but also challenging for clustering analysis and gene network inference from scRNA-seq data. Most existing single-cell clustering algorithms select highly variable genes first and then perform principal component analysis (PCA) dimension reduction followed by graph-based or consensus k-means clustering^5, 8^. The selection of top variable features improves the clustering speed but is arbitrary and may lose the information that can distinguish close cell states. Furthermore, the linear-transformation of PCA and co-expression analysis using linear Pearson or Spearman correlations^6^ may not capture the nonlinear cell-cell distance and gene-gene correlations.

To address the above challenges, we developed a mutual information (MI)-based integrative computational framework, termed single-cell Mutual Information-based Network Engineering Ranger (scMINER). ScMINER was designed to perform unsupervised clustering and reverse engineering of cell-type specific TF and SIG networks from scRNA-seq data. For this, we leveraged SJARACNe^9^, an MI-based algorithm for gene network reconstruction from bulk omics data^13^, to infer cluster-specific TF and SIG networks from scRNA-seq profiles. Based on the data-driven and cell-type specific networks, scMINER was able to transform the single-cell gene expression matrix into single-cell activity profiles and then identify cluster-specific TF and SIG drivers including hidden ones that show changes at the activity but not expression level. We benchmarked the clustering performance of scMINER in 11 scRNA-seq datasets against three widely-used tools (Seurat^10^, SC3^11^, and Scanpy^12^), and showed that scMINER outperforms the other methods. In particular, scMINER improves the separation of similar cell types, thus significantly increasing the signal-to-noise-ratio for downstream cell-type specific network reconstruction and hidden driver identification. We demonstrated the power of scMINER in single-cell studies of immune cell diversity in peripheral blood mononuclear cells (PBMCs), exhausted T cell lineage differentiation, and tissue specification of regulatory T (Treg) cells.

## Results

### Overview of scMINER

To characterize nonlinear relationships among cells and genes from single-cell omics data, we developed a MI-based scMINER workflow. In scMINER, we used nonlinear MI to measure cell-cell similarity for unsupervised clustering analysis and gene-gene correlation for reverse-engineering of cluster-specific intracellular networks from scRNA-seq data, which enables the identification of cell type-specific hidden drivers and their network rewiring events. Specifically, scMINER was designed to include two key components (**Fig. 1**): (i) Mutual Information-based Clustering Analysis (MICA) and (ii) Mutual Information-based Network Inference Engine (MINIE). We chose MICA because it uses MI to quantify cell-cell distance, which allows for

**Figure 1:**
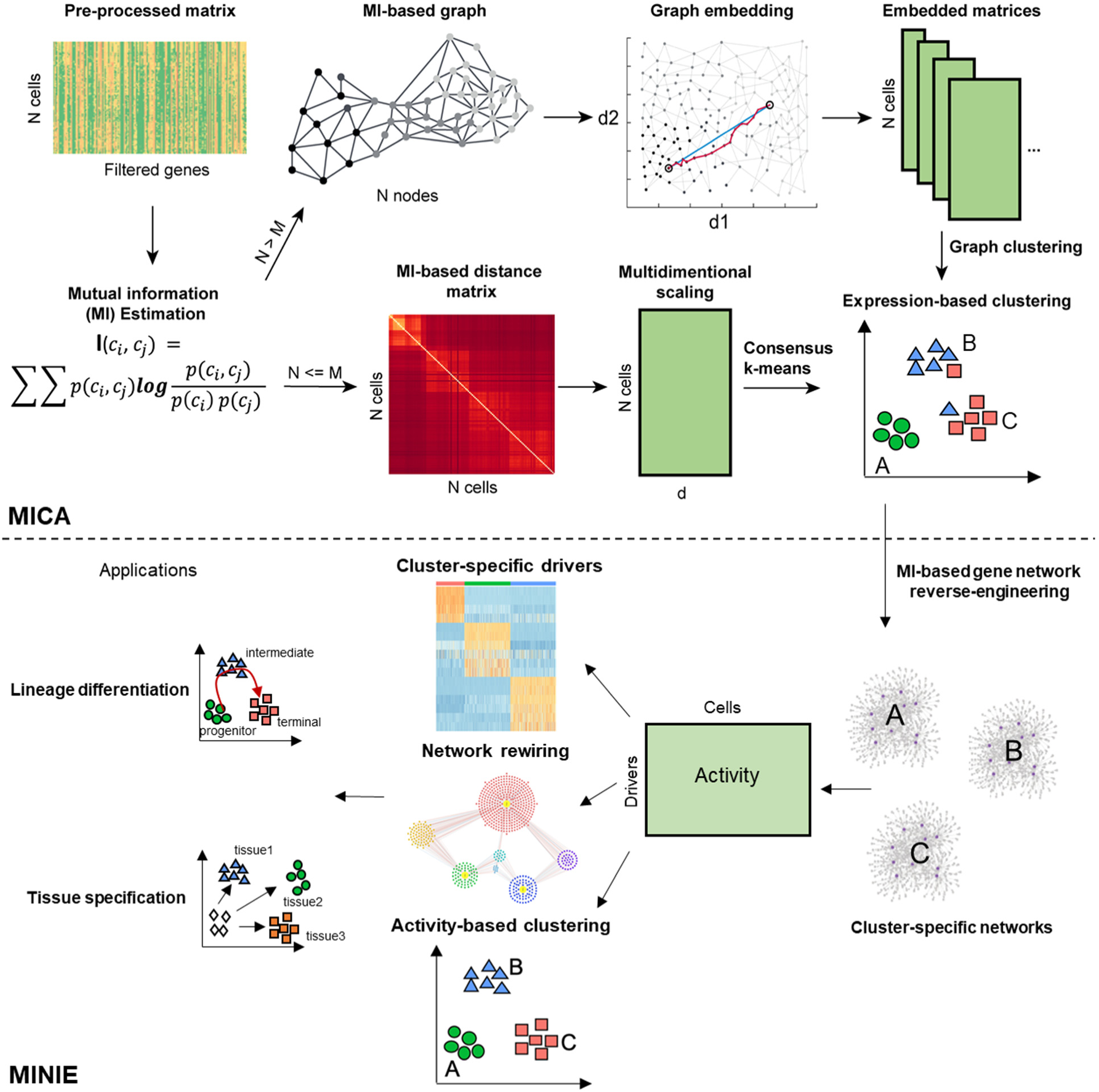
Overview of scMINER. scMINER is a system biology toolkit that has been separated into mutual information-based clustering analysis (MICA) and mutual information-based network inference engine (MINIE). Mutual information (MI) is first calculated from the gene count matrix from scRNA-seq to obtain a MI-based distance matrix. Multidimensional scaling (MDS) based dimension reduction is then performed followed by K-mean clustering. Cell state specific networks are constructed using SJARACNE to infer the regulons of transcription (TF) and signaling protein (SIG) drivers. The importance of each TF and SIG driver is measured by comparing between different cell states. The regulatory network re-wiring of these drivers in various cell states could be further captured, which serves the basis of identifying critical drivers in cell lineage differentiation and tissue specific specification. Moreover, the regulon activity could be further used to refine expression-based clustering (activity-based clustering). characterization of the intrinsic nonlinear similarity of gene expression distributions among cells. To balance the efficiency and accuracy of clustering, we implemented MICA that combines two prevalent strategies for clustering analysis of scRNA-seq data^5, 8^: a graph-based approach (e.g., Seurat^10^, Scanpy^12^), which is fast and builds a heuristic cell-cell graph and applies community detection, and consensus k-means-based clustering (e.g., SC3^11^), which is slower but more accurate, as it iteratively identifies the globally optimal k clusters and uses a consensus approach to increase the robustness. Thus, we took advantage of both strategies to balance the clustering stability and scalability, resulting in MICA that uses graph-based clustering when the cell number is large (default 5,000) and uses the k-means-based approach otherwise. We also employed nonlinear graph embedding (GE) in graph-based clustering and multidimensional scaling (MDS) in k-means-based clustering to reduce the noise arising from the intrinsic “dropout” effects in scRNA-seq data, as well as optimization of the number of dimensions used for clustering.

MICA was integrated with MINIE that uses clustering results to reverse-engineer intracellular gene networks for each of the clusters by using a modified MI-based algorithm, SJARACNe^9^. Although originally developed to analyze bulk omics data, we re-parameterized SJARACNe to handle single-cell transcriptomics data. To overcome the sparseness of scRNA-seq data for gene network inference, we designed MINIE to employ a MetaCell^13^ approach by aggregating gene expression profiles of similar cells to reconstruct cluster-specific TF and SIG networks from scRNA-seq data. Furthermore, for each TF or SIG candidate driver, MINIE infers the cluster-specific gene activity based on the expression of its predicted regulon genes in the corresponding cluster. Taken together, the combination of non-linear nature of MICA clustering, and the activity-based analysis performed by MINIE is expected to overcome the dropout effects of scRNA-seq data and identify cluster-specific hidden drivers. Therefore, we propose that scMINER represents a robust platform for identification of cluster-specific hidden drivers and their target rewiring in lineage differentiation, tissue specification and many other biological processes.

### scMINER outperforms popular single-cell clustering tools

To benchmark the clustering performance of scMINER, we considered the three widely-used methods: Seurat^10^ and Scanpy^12^, representing graph-based approaches, and SC3^11^ representing k-means-based algorithms. These methods were also among the top performers based on the previous evaluations^14^. We used 11 scRNA-seq datasets from different platforms with known cell-type labels and with various numbers of cells (**Supplementary Table 1**). These datasets consist of four gold-standard and three silver-standard datasets used for benchmarking SC3, as well as four additional large datasets with cell-type labels based on cell sorting markers and expert curations^15, 16^. We used the Hubert-Arabie Adjusted Rand index (ARI), which ranges from 0 for random to 1 for identical matching, to quantify how well the inferred clusters by different methods recovered the reference labels.

The benchmarking analysis indicated that MICA (in scMINER) consistently outperformed the other three methods with the highest ARI across all the datasets (**Fig. 2a, Supplementary Fig. 1a**), except for the Klein dataset, where the ARI of MICA, Seurat, and SC3 were almost identical and significantly higher than the ARI of Scanpy. Among the three benchmarking algorithms, there was no consistent winner: SC3 significantly outperformed graph-based Seurat and Scanpy in a few datasets with a small number of cells (e.g., Buettner and Pollen datasets), but failed for the two datasets with a large number of cells (e.g., Zheng and Bakke datasets); Seurat performed well in a few small datasets (e.g., Yan and Goolam datasets). Overall, MICA is the most consistent method with an average ARI value of 0.83 and the lowest variance (**Fig. 2b**). The superior performance of MICA was also confirmed by using an alternative metric, the Adjusted Mutual Information (AMI; **Supplementary Fig. 1b**).

**Figure 2:**
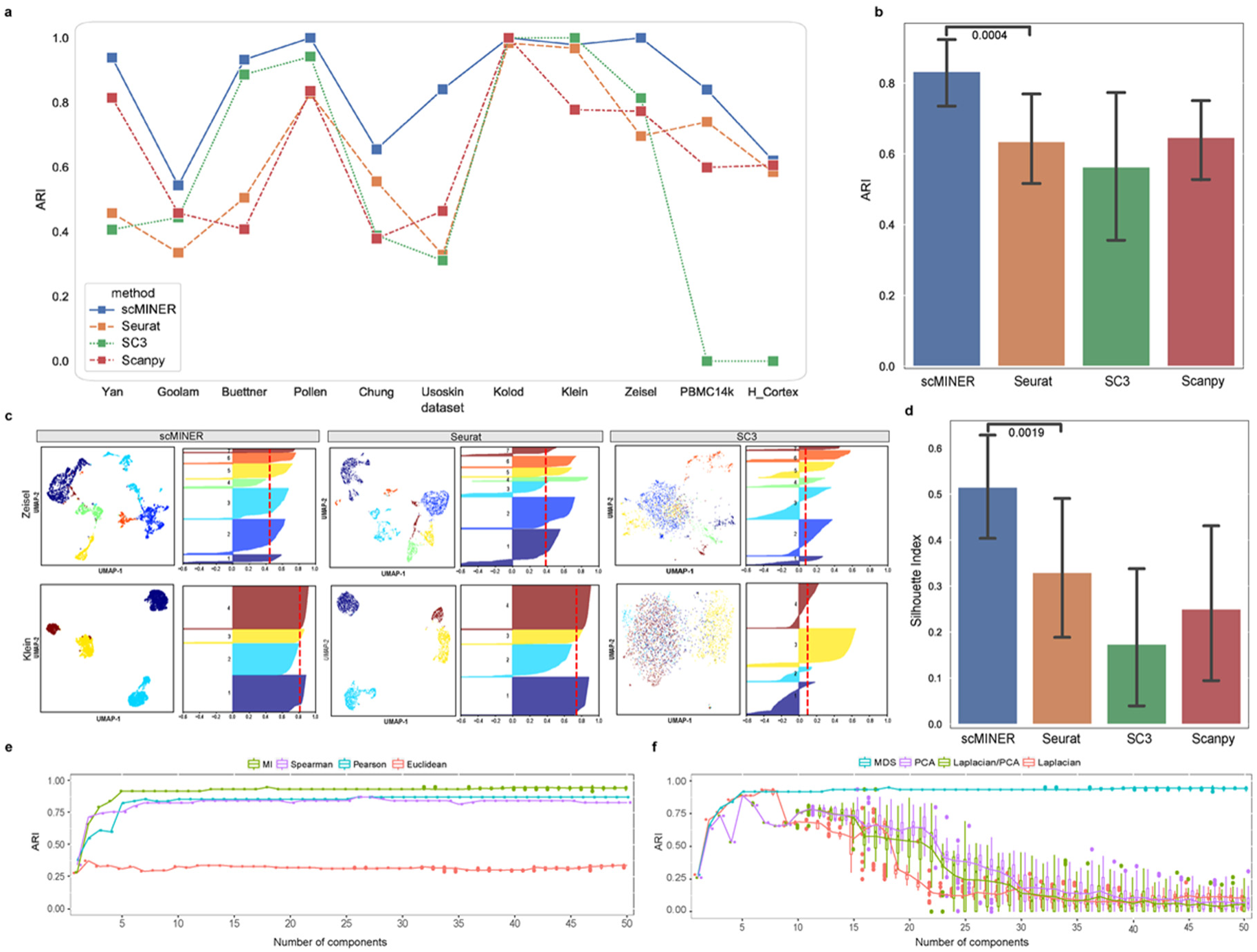
Evaluation of scMINER clustering performance. **a,** Clustering performance of scMINER, Seurat, SC3 and Scanpy measured by adjusted Rand index (ARI). **b,** The average ARIs and their variance (vertical segments). scMINER significantly outperforms other clustering methods (*p* = 0.0004 by the one-sided Wilcoxon test). **c,** UMAP and silhouette plots of the Zeisel and Klein datasets using scMINER, Seurat, and SC3. Silhouette index is reported (red dashed line) for each UMAP representation of clustering result. **d,** Average silhouette index values and their variance (vertical lines). scMINER significantly outperforms other clustering methods (*p* = 0.0019 by the one-sided Wilcoxon test). **e,** Clustering performance comparison using four distance metrics mutual information (MI), Spearman correlation, Pearson correlation and Euclidean distance as metrics. MI outperforms other linear metrics when the number of dimensions is greater than a fixed number. **f,** Clustering performance comparison using four dimension reduction approaches multidimensional scaling (MDS), principal component analysis (PCA), Laplacian, and PCA sequentially followed by Laplacian (LPCA).

To evaluate MICA’s effect on visualization via UMAP, we calculated the silhouette index (SI)^17^ scores of 2d-UMAP visualization against the reference labels for each clustering method in all 11 benchmarking datasets. We observed that MICA exhibited much higher SI values than Seurat, SC3, and Scanpy (**Fig. 2c**), suggesting higher purity and closer to true biological cell types of MICA clustering than those of clustering by the other methods. Again, MICA is the most consistent method as shown by the lowest variance across all the datasets (**Fig. 2d**).

We reasoned that the consistent superior clustering performance of MICA was because of the use of nonlinear metrics, including MI for quantifying cell-cell similarity and GE for dimension reduction. To investigate this further, we focused on the four gold-standard data sets with ground-truth data (Yan, Pollen, Kolodziejczyk, and Buettner datasets). First, we replaced the default MI metric with other widely-used distance metrics including Euclidean, Pearson, and Spearman correlation coefficients while keeping other steps of the MICA workflow the same. For each distance metric, we performed the clustering on the four gold-standard datasets for a range of MDS components from 1 to 50 (**Fig. 2e, Supplementary Fig. 2a**). MI-based clustering achieved consistently better performance than the other three metrics regardless of the number of components, which indicates that MI is more robust in capturing the actual cell-cell similarity, likely due to its non-linear nature.

Similarly, we benchmarked MDS with three other commonly used dimension reduction methods: 1) Principal Component Analysis (PCA), 2) Laplacian, 3) Parallel PCA and Laplacian (Laplacian/PCA) by changing the dimension reduction metric only in the MICA-k-means pipeline. Clustering accuracy (ARI) was measured with the increment of selected components after reduction, ranging from the 1st to 50th, with downstream steps remaining the same. The results from the four gold-standard datasets showed that MDS reached its maximum accuracy during the process and remained stable with addition of more components, while the other three approaches reached their respective best ARI at different numbers of components and failed to maintain the optimal result when more components were involved (**Fig. 2f, Supplementary Fig. 2b**). This behavior of MDS enabled us to optimize the selection of components, a critical parameter for downstream k-means clustering, and we set 19 as the default in the MICA-MDS pipeline as this number ensured we reached the maximum accuracy in all datasets we have benchmarked.

We also evaluated the computing resource usage of MICA-GE and its distribution in each step using the two largest Zheng and Bakken datasets (**Supplementary Fig. 3a**). The MI-kNN step used ∼60% of the total time. We also evaluated the effects of GE parameters (e.g., number of workers, number of kNN neighbors, node2vec window size, etc) on clustering performance (**Supplementary Fig. 3b**), which helped to optimize the parameters. Taken together, scMINER-MICA is a robust, accurate, and efficient clustering algorithm.

### scMINER improves the clustering of ambiguous T-cell subpopulations in PBMCs

To document the intrinsic clustering performance on a well-characterized mixed cell population, we compiled a training dataset with 14,000 PBMCs forming seven mutually exclusive cell types from the Zheng dataset^15^. We picked clustering resolution parameters for MICA and Seurat to produce the number of clusters based on the known number of cell types, and examined the true labels of the cells as defined by fluorescence-activated cell sorting (FACS) (**Fig. 3a**). Though CD4^+^ and CD8^+^ differences were well-recaptured by both MICA and Seurat, we found dramatic differences in identifying CD4^+^ cell subpopulations by MICA and Seurat (**Fig. 3b, c**). Specifically, MICA clearly identified a central memory T-cell population (CD4^+^/CD45RO^+^ memory T) and a Treg population (CD4^+^/CD25^+^ regulatory T), whereas Seurat generated two clusters (cluster 1 and 2) of CD4^+^ cells with mixed subpopulations (53% of CD4^+^ central memory cells and 45% of CD4^+^ Tregs, **Fig. 3d**). MICA identified three subpopulations of CD4^+^ T cells and one subpopulation of CD8^+^ naive cytotoxic T cells. In contrast, Seurat failed to separate the subpopulations of CD4^+^ T cells even after tuning the resolution parameter to produce 8 clusters. To avoid over-emphasizing the ground-truth labels which arises from the one or two surface-markers, we compiled a set of cell type specific signature genes and calculated the signature scores for each of the MICA and Seurat clusters (**Fig. 3c**). The signature scores show that Seurat identified two monocyte clusters and failed to identify pure clusters of CD4^+^ central memory T cells and CD4^+^ Tregs. Further, by comparing the two presumably Treg clusters, cluster 2 in MICA and cluster 1 in Seurat, we found that the number of cells with well-known Treg markers (FOXP3, IL2RA, TIGIT)^18, 19^ in the MICA cluster is much higher than the Seurat cluster (**Fig. 3e**). The high signal-to-noise ratio highlights the advantage of MICA over Seurat to uncover cluster-specific signals for downstream network analysis. Additionally, we found that the high purity and the matching of the number of clusters to that of cell types are partially due to the amplified signals by MICA’s default count-per-million reads normalization approach (**Fig. 3f, Supplementary Fig. 4).**

**Figure 3:**
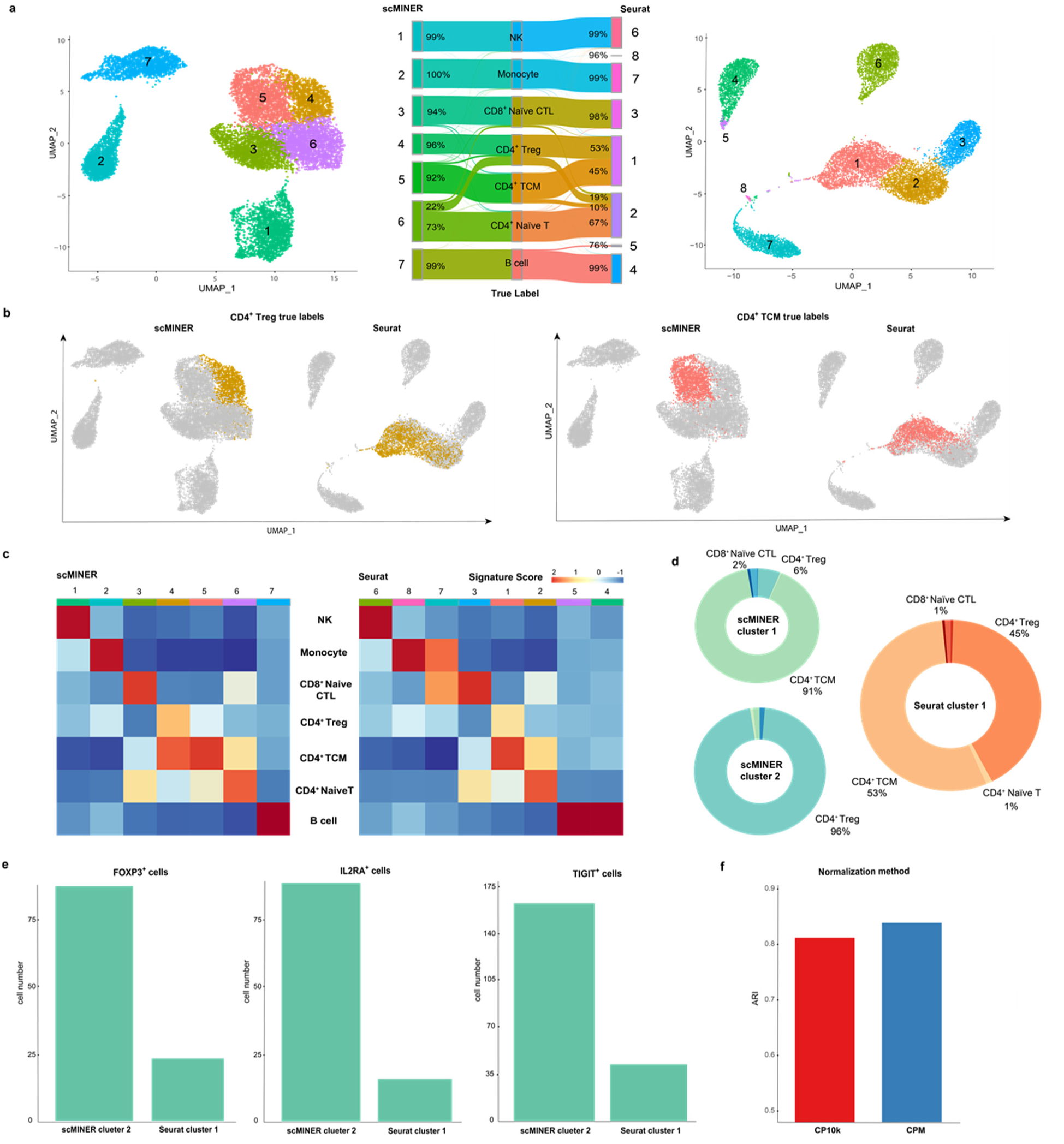
scMINER improves the clustering of ambiguous subpopulations in PBMCs in comparison with Seurat. **a,** UMAP and Sankey plots of scMINER and Seurat clustering results annotated using true labels. **b,** CD4TCM and CD4Treg cells (by true label) projected on scMINER and Seurat clustering UMAP plots. **c,** Signature score heatmap plots of scMINER and Seurat clusters calculated using curated markers for each cell type. **d,** Donut plots of scMINER cluster 1, 2 and Seurat cluster 1 to show the cell type compositions and the cluster purity. **e,** Comparison of the number of CD4Treg cells expressing master regulators FOXP3, IL2RA, and TIGIT on scMINER cluster 2 and Seurat cluster 1. **f,** MICA clustering performance comparison using CPM and CP10k normalization methods.

Every clustering approach has a few intrinsic parameters that could potentially change the clustering results. Therefore, it is instructive to examine the robustness of these parameters, such as the resolution parameter in the Louvain algorithm. We used CD4^+^ Tregs as a proxy to examine how the clusters vary across different resolutions and cluster counts (**Supplementary Fig. 5**). The analysis revealed that CD4^+^ Tregs spread across more Seurat clusters with increasing resolution, whereas most CD4^+^ Tregs form a single MICA cluster regardless of the resolution. Given that Seurat employs a variation-based gene selection step before PCA analysis, we examined whether the number of highly variable genes impacts the clustering performance and so no improvement on Seurat’s ability to characterize CD4^+^ Treg cell similarities (**Supplementary Fig. 6**). Taken together, these studies indicate that scMINER achieves improved clustering of ambiguous cellular subpopulations.

### scMINER infers immune marker protein activity and improves clustering of PBMCs

With improved clustering, we then applied the scMINER-MINIE workflow to map cell-type-specific intracellular TF and SIG networks from scRNA-seq data. Therefore, this process allowed us to transform the single-cell expression profiles into single-cell protein activity profiles, and identify hidden drivers underlying each cell type that expression may fail to capture. We continued the analysis of the PBMC dataset with 7 cell types – monocytes, B cells, NK cells, CD4^+^ T cells, and naïve CD8^+^ T cells (**Fig. 4a**). With MINIE, we first generated cell-type-specific TF and SIG networks based on scRNA-seq profiles of each of the seven sorted populations containing 2,000 cells (**Supplementary Fig. 7a**). We then inferred TF and SIG activities by taking the expression level of predicted targets into account, resulting in an activity matrix of 1,428 TFs and 3,382 SIGs. The activity matrix overcame the sparseness of scRNA-seq data; it satisfied a normal distribution, which drastically cut down the theoretical pre-requisite for any statistical testing.

**Figure 4:**
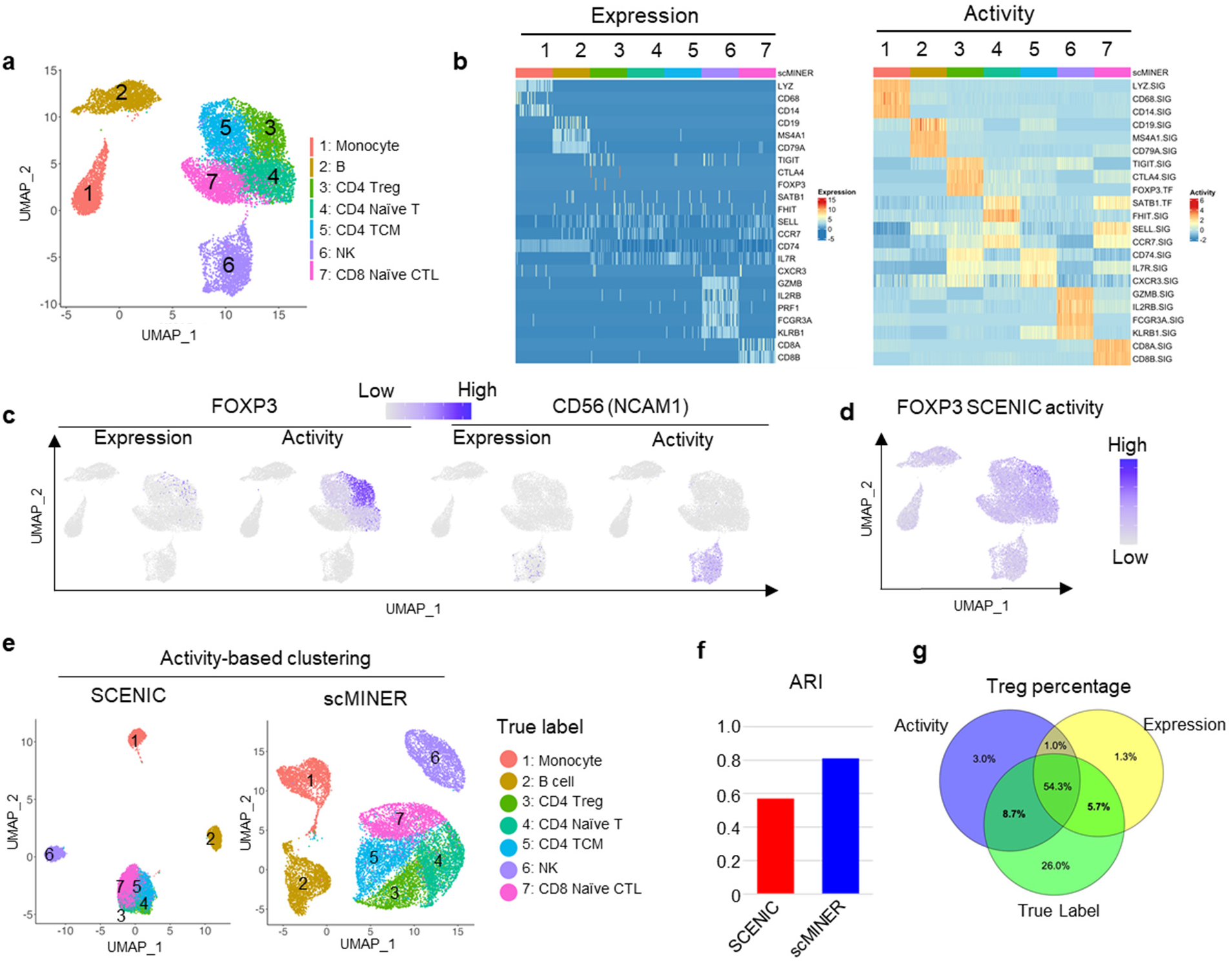
Comparison of scMINER and SCENIC on activity-based mark identification and clustering of PBMC cell types. **a,** Unsupervised scMINER clustering of 7 clusters of sorted PBMC cell types on UMAP. **b,** Heatmap visualization of cell marker expression (left) and predicted activity (right) in each cell from sorted PBMC cell types. The regulatory networks are generated in cell-type specific manner. **c,** FOXP3 (left) and CD56 (encoded by NCAM1, right) expression and scMINER activity on UMAP. **d,** UMAP visualization of FOXP3 activity predicted by SCENIC pipeline. **e,** Unsupervised clustering of the 7 sorted PBMC cell types based on SCENIC and scMINER activity. The true labels of these 7 cell types are labeled. **f,** The Adjusted Rand Index (ARI) of clustering in **e** based on SCENIC and scMINER activity.

Unique molecular signatures have been used in literature to define different immune cells in PBMCs, including monocytes (LYZ, CD68, and CD14), B cells (CD19, MS4A1, and CD79A), Treg cells (CTLA4, TIGIT, and FOXP3), NK cells (GZMB, IL2RB, FCGR3A, and KLRB1) and naïve CD8^+^ T cells (CD8A, CD8B, SELL)^20–22^. However, the expression level of a few markers cannot identify and separate immune cell subtypes, especially CD4^+^ T cell subsets in this PBMC dataset, possibly due to gene dropout (**Fig. 4b**). Compared to expression, the activity of classical immune markers can well separate these immune cell types (**Fig. 4b**). The dropout of both Treg marker FOXP3 and NK cell marker CD56 (encoded by NCAM1 gene) marker expression can be rescued by their effect on the UMAP (**Fig. 4c**), as well as other cell type-specific markers such as CD19, CD8A, and CD14 (**Supplementary Fig. 7b**). The activity of their signatures further separated the subsets of CD4^+^ T cells. For example, naïve CD4^+^ T cells showed higher FHIT and SATB1 activity and lower CXCR3 activity than CD4^+^ memory T cell subsets (**Fig. 4b and Supplementary Fig. 7c**). Compared with the widely used scRNA-seq regulon analysis method SCENIC (**Fig. 4d**), scMINER was able to identify Treg cells with FOXP3 activity with more specificity (**Fig. 4c**).

The activity calculated from scMINER could also be used to improve clustering unbiasedly. We calculated the activity of the cell type signatures based on GRN generated from MetaCell^13^ using the total PBMC cells (**Fig. 4f**). We observed that MINIE-based activity clustering outperformed SCENIC-based activity clustering, in terms of recovering the reference labels. The separation of Treg cells from other T cells was further improved compared to gene expression-based clustering (**Fig. 4g**). Taken together, scMINER-inferred activity overcomes the “dropout” effects to improve marker protein identification and clustering from scRNA-seq data.

### scMINER reveals drivers and their network rewiring in exhausted CD8^+^ T cell differentiation

Next, we demonstrated the power of scMINER to identify drivers in cell lineage and differentiation using a specific example of exhausted CD8^+^ T cell differentiation, a phenotype associated with severe infection, cancer and autoimmunity^23–26^. Previous studies have shown that the exhausted CD8^+^ T cells contain heterogeneous subpopulations with differential capability to respond to anti-PD1 therapy^27–33^. We performed scMINER clustering on scRNA-seq profiles of exhausted CD8^+^ T cells in mice chronically infected with lymphocytic choriomeningitis virus (LCMV) Clone 13 (Cl13) at a late stage (day 28)^32^. We recapitulated the 3 major subpopulations of exhausted CD8^+^ T cells defined by TCF-1 (encoded by *Tcf7* gene), CX3CR1, and TIM3 (encoded by *Hacvr2*)^+^: TCF-1^+^ exhaustion progenitor (Tpex), CX3CR1^+^ effector-like (Teff-like, effector-like Tex), and CX3CR1^−^TIM3^+^ terminal exhausted T cells (Tex) (**Fig. 5a**). These three populations are distinct both phenotypically and functionally^32, 34, 35^.

**Figure 5:**
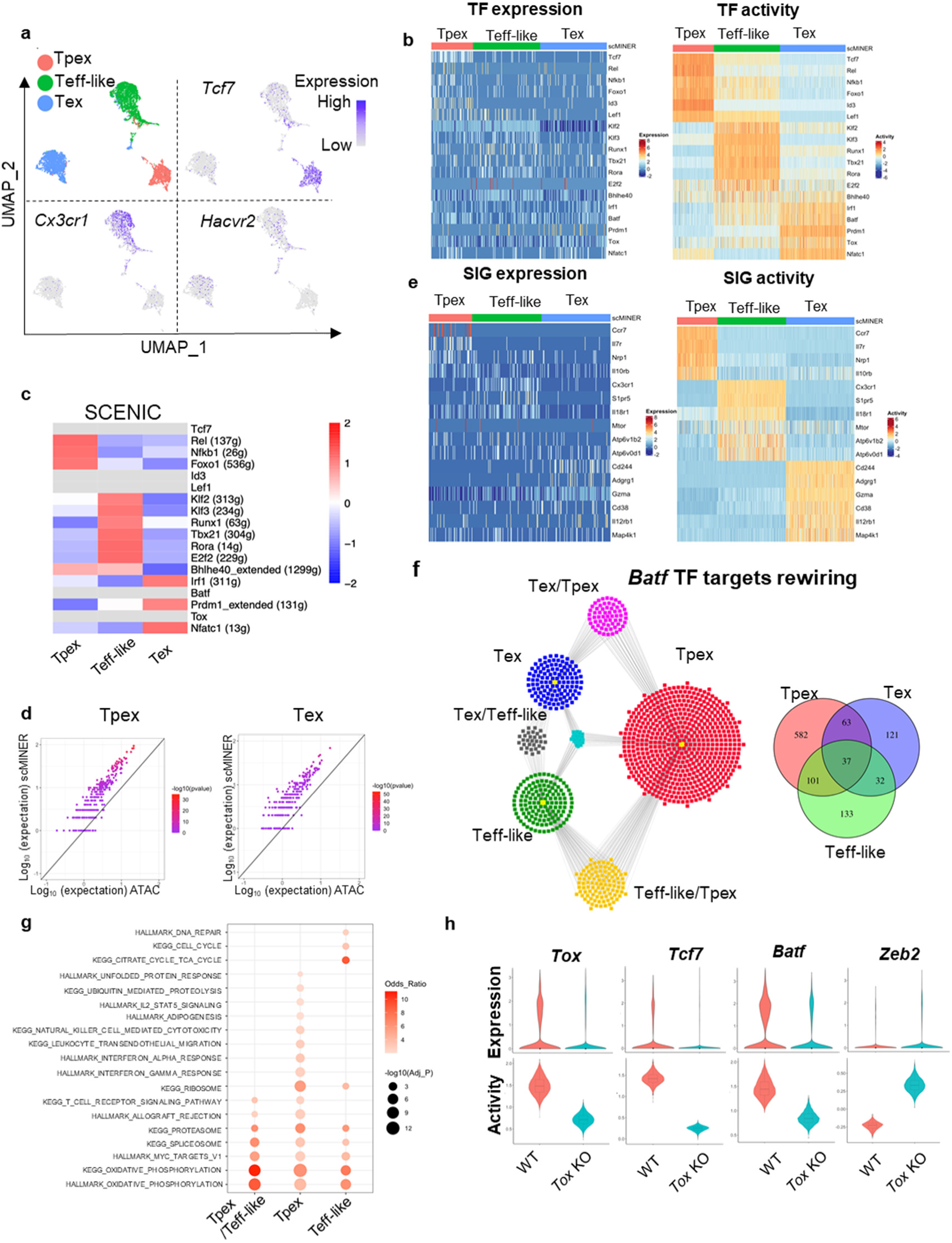
scMINER captures cluster-specific drivers for CD8^+^ exhausted T cells. **a,** MICA MDS clustering of GP33-tetramer^+^ CD8^+^ T cells at day 28 from mice chronically infected with LCMV Clone 13 (GSE122712). The expression of *Tcf7*, *Cx3cr1* and *Havcr2* is visualized on UMAP. **b,** Heatmap visualization of expression (left) and predicted activity (right) of selected TFs in each cell from 3 subsets of CD8^+^ T cells. **c,** Heatmap visualization of the average TF SCENIC activity in each cluster of CD8^+^ T cells. TFs without activity predicted by SCENIC are shown in light grey. **d,** Similarity of TF regulon in Tpex and Tex CD8^+^ T cells generated by SJARACNe and footprint genes detected by ATAC-seq data (GSE123236) in corresponding cell clusters. Expected number of genes in intersection of ATAC-seq footprints as reference (log10 scale, x axis) with regard to hypergeometric distribution vs. observed intersection (log10 scale, y axis). For all genes, the observed intersection is significantly higher than expectation (black line). The color of the dots represents −log10 (P-value) according to Fisher’s exact test. **e,** Heatmap visualization of cell marker expression (left) and predicted activity (right) of selected SIGs in each cell from 3 subsets of CD8^+^ T cells. **f,** The regulons of Batf in Tpex (red), Teff-like (green) and Tex (blue) cells. Regulons shared by 2 or more cell types are highlighted as yellow (Tpex and Teff-like), grey (Teff-like and Tex), magenta (Tex and Tpex), and turquoise (Tpex, Teff-like and Tex). **g,** Functional pathway enrichment of Batf regulons of Tpex and Teff-like cells and the one shared by Tpex and Teff-like. **h**, Violin plot of the expression (upper) and activity (lower) of Tox, Tcf7 and Batf in wild-type and Tox deficient CD8^+^ T cells.

We then applied scMINER to delineate the transcription regulatory networks and underlying hidden drivers along the state changes from Tpex to Teff-like to Tex. Previous studies reported that TCF-1, FOXO1, ID3, LEF1, and NF-kB related transcriptional factors REL and NFKB1 are involved in Tpex regulation^36, 37^, although the expression levels of these TFs didn’t show marked changes (**Fig. 5b**). In contrast, scMINER-inferred activities of these TF regulators based on cluster-specific regulatory networks uncovered their importance in Tpex (**Fig. 5b**). The surprising enrichment of NF-kB-related transcriptional factors suggested possible roles of NF-κB signaling pathway in regulating the formation or maintenance of Tpex cells. For the Teff-like subpopulation, T-bet (encoded by *Tbx21*) is known to be important for the effector function of this population^36^ and indeed has heightened scMINER activity (**Fig. 5b and Supplemental Fig. 8a**). Other TFs such as KLF2/3, RUNX1, and ROR-α (encoded by *Rora*) also have increased regulon activities in the Teff-like cluster, consistent with the prediction by SCENIC analysis in the literature^36^. Finally, the terminally exhausted Tex cells showed increased activity of NFATC1, BLIMP1 (encoded by *Prdm1*), and TOX, which were reported to promote exhaustion^36, 38–41^. Notably, the increased scMINER activities for T-bet and BLIMP1 in Teff-like and Tex respectively are much more obvious than their expression. Although BATF function in terminal exhaustion and effector function is still debatable depending on the biological context^36, 42–45^, overexpressed BATF has been shown to be critical for regulating effector function in adoptively transferred antigen-specific CD8^+^ T cells and CAR-T cells^45, 46^. The scMINER analysis was able to accurately detect BATF activity that was missed by analysis of its expression pattern alone (**Supplementary Fig. 8a**).

The TF transcriptional networks of these three CD8^+^ T cell exhaustion stages also captured known master TF drivers in CD8^+^ T cells. For example, in Tpex cells, the TCF-1 is predicted to directly promote expression of *Cd9* and *Ms4a4c* (**Supplementary Fig. 8b**), while FOXO1 is known to promote *Pdcd1* expression during exhaustion^36, 47^. In Teff-like cells, T-bet regulates *Zeb2* expression, which has been reported to be critical for effector CD8^+^ cytotoxic cell differentiation^48, 49^. In terminal Tex cells, TOX regulates co-inhibitory molecular *Lag3* expression^50^.

To benchmark the performance of scMINER, we compared it with SCENIC^6^. SCENIC was applied to the exhausted CD8^+^ T cell dataset and was able to capture a few positive control drivers of T cell exhaustion, including REL, NFKB1, FOXO1, KLF2/3, RUNX1, T-bet, ROR-α, and E2F2, all of which were predicted by scMINER (**Fig. 5c**). However, SCENIC failed to predict TCF-1, ID3, LEF1, BATF, and TOX activity in this dataset based on their co-expressed regulons (filtered out in motif2tf step), among which TCF-1, BATF, and TOX are well-established TF regulators of exhausted T cell differentiation. This suggested that scMINER goes beyond the limitation of TF motif database and can predict a broader spectrum of TF drivers in exhausted T cell differentiation directly from scRNA-seq data. Intriguingly, we also found the TF regulons predicted by scMINER have a significant overlap with TF footprint genes detected by ATAC-seq analysis^51^ of Tpex and Tex cells. This suggested that scMINER regulons reflect true transcriptional targets of TFs in a context-specific fashion (**Fig. 5d**).

Distinct from SCENIC that is focused on TF driver inference, scMINER can be used to reconstruct context-specific signaling networks and infer the activity of signaling factors. Indeed, scMINER successfully identified that memory-associated surface markers such as CCR7 and IL7R are uniquely activated in Tpex cells (**Fig. 5e**). CX3CR1, a surface hallmark for Teff-like cells, also exhibited higher activity in Teff-like cells. Effector function of CD8^+^ T cells is correlated with high mTOR activity^34, 52^, and scMINER did capture that the activities of mTOR and its upstream regulator V-ATPases^53^ were high in Teff-like cells (**Fig. 5e and Supplementary Fig. 8c**). Further, genes that were reported to be highly expressed in terminal Tex cells such as *Cd244* (2B4)^54^ and *Cd38*^55^ exhibited the highest activity in Tex cluster (**Fig. 5e**). MAP4K1 (encoded by *Hpk1*) is known to promote terminal exhaustion^56^, and scMINER captured the most increased activity in terminal Tex cells. Notably, the selective kinase activity of mTOR and MAP4K1 in Teff-like and Tex could not be captured by their gene expression changes from scRNA-seq data (**Fig. 5e**). All of the above indicates that scMINER can capture context-dependent activity of signaling proteins, including surface receptors, intracellular enzymes, and kinases.

By inferring cell-type-specific networks for various clusters, scMINER can uncover the regulon rewiring of drivers among cell types, and thus determine the transcription regulation during cell state transition. BATF has recently been identified to have a role in promoting Tpex to Teff-like transition via different regulons^36^. Indeed, scMINER network analysis captured that BATF regulon targets were significantly rewired among Tpex, effector-like Tex, and terminal Tex states (**Fig. 5f**). In the overlapped BATF regulons between Tpex and effector-like Tex, BATF regulates effector CD8^+^ T cell-associated oxidative phosphorylation and T cell receptor signaling pathway (**Fig. 5g**), which suggested BATF may promote metabolic rewiring to increase effector-like Tex and is consistent with the role of BATF reported in the literature^45, 46^. In contrast, BATF regulons that are unique in Tpex cells were enriched in cytotoxic IFN-α and IFN-γ response pathways while BATF regulons that are unique in Teff-like cells were enriched in cell cycle-related pathways (**Fig. 5g**). Together, these results indicated that BATF may alter its regulons in different cell subsets to tune their activity and exert their regulatory function during the process of CD8^+^ T cell exhaustion.

Understanding complex transcriptional network changes in response to gene perturbation is among the biggest challenges for mechanism studies. Since TOX is a master TF regulator of T cell exhaustion, we analyzed scRNA-seq data (GSE119940) profiling wild-type (WT) and TOX knockout (KO) CD8^+^ T cells^38^ during chronic infection to examine whether scMINER can reveal the regulatory circuits of TOX in chronic infection (**Supplementary Fig. 8d**). TOX-KO CD8^+^ T cells exhibited reduced TOX activity compared to WT cells (**Fig. 5h**), indicating scMINER can correctly capture TOX activity in this biological context. *Tcf7* expression was reported to be downregulated upon TOX KO^38, 39^, and we indeed found that TOX KO decreased both expression and activity of *Tcf7* (**Fig. 5h**). Since *Tcf7* marks the Tex cell precursors (Tpex), our results aligned with previous reports that the primary defect in TOX-KO exhausted T cells was the inability to rewire the transcriptional control of *Tcf7* after the initial development of Tpex. Apart from the reduction of *Tcf7* activity, TOX-KO CD8^+^ T cells were also accompanied by the reduced activity of BATF (**Fig. 5h**), which is required for sustaining antiviral CD8^+^ response during chronic infection^57^. TF motif enrichment analysis also validated the reduced activity of BATF in TOX-KO CD8^+^ T cells (**Supplementary Fig. 8e**). Moreover, the TOX-KO cells also increased the activity of effector-related transcription factor ZEB2 (**Fig. 5h**), in line with the increased effector T cell function in TOX-KO cells, which cannot be solely explained by loss of *Tcf7*^39^. Moreover, other top TF and SIG drivers predicted by scMINER in TOX-KO CD8^+^ T cells were also enriched in the effector T cell signature but not exhausted T cell signature (**Supplementary Fig. 8f**). These results together highlighted that a complex transcription regulatory network, not just *Tcf7* expression, is required for TOX-mediated effector function and exhaustion progression in CD8^+^ T cells during chronic infection.

### scMINER exposes drivers underlying Treg tissue specification

While different types of T cells play different roles, the same type of T cell could have specific roles in specific tissues, driven by the underlying transcriptional regulatory networks^58^. For example, tissue specific Treg cells not only maintain immune tolerance but also promote homeostasis and regeneration after tissue damage^59, 60^. To examine this further, we used scMINER to dissect TF and SIG drivers underlying tissue specification of Treg cells. We performed scMINER clustering of scRNA-seq profiles of Treg cells from different tissues including spleen, lung, skin and visceral adipose tissue (VAT)^61^. Clustering results exhibited that Treg cells were well separated by their tissue origins. Skin and VAT Treg cells displayed high *Cd44* and low *Sell* expression, while spleen and lung Treg cells showed the opposite (**Fig. 6a**). We then reconstructed tissue-specific regulatory networks of Treg cells to establish the regulons of TF drivers for their activity inference. Differential activity analysis by scMINER revealed that resting Treg cell-associated TFs (e.g., TCF-1, KLF2, SATB1, and BACH2) all have the highest activity in spleen and lung Treg cells (**Fig. 6b and c**). Endoplasmic reticulum (ER) stress is critical for normal skin function and associated with skin-related autoimmune disease^62^. Interestingly, Treg cells in the skin upregulate the activity of ER stress regulator ATF6, suggesting a possible influence from the skin microenvironment (**Fig. 6b and c**). VAT Treg is the most well-characterized tissue Treg with high activity of PPARG, FLI1, RORA, and RARA, consistent with the prediction based on motif enrichment analysis of ATAC-seq data^63^ and literature^61^ (**Fig. 6b right**). Notably, scMINER activity-based TF driver inference captured these tissue-specific drivers, which could not be clearly identified from their expression patterns (**Fig. 6b left**). Orthogonal scATAC-seq-based TF accessibility analysis (**Fig. 6c**) and SCENIC analysis (**Supplementary Fig. 9a**) captured differential tissue-selective activity of BACH2, KLF2, ATF6, and PPARG, but their signals were much weaker than scMINER-based activity analysis, and could not correctly predict the high FLI1, RORA, and RARA activity in VAT Treg cells (**Supplementary Fig. 9b**).

**Figure 6:**
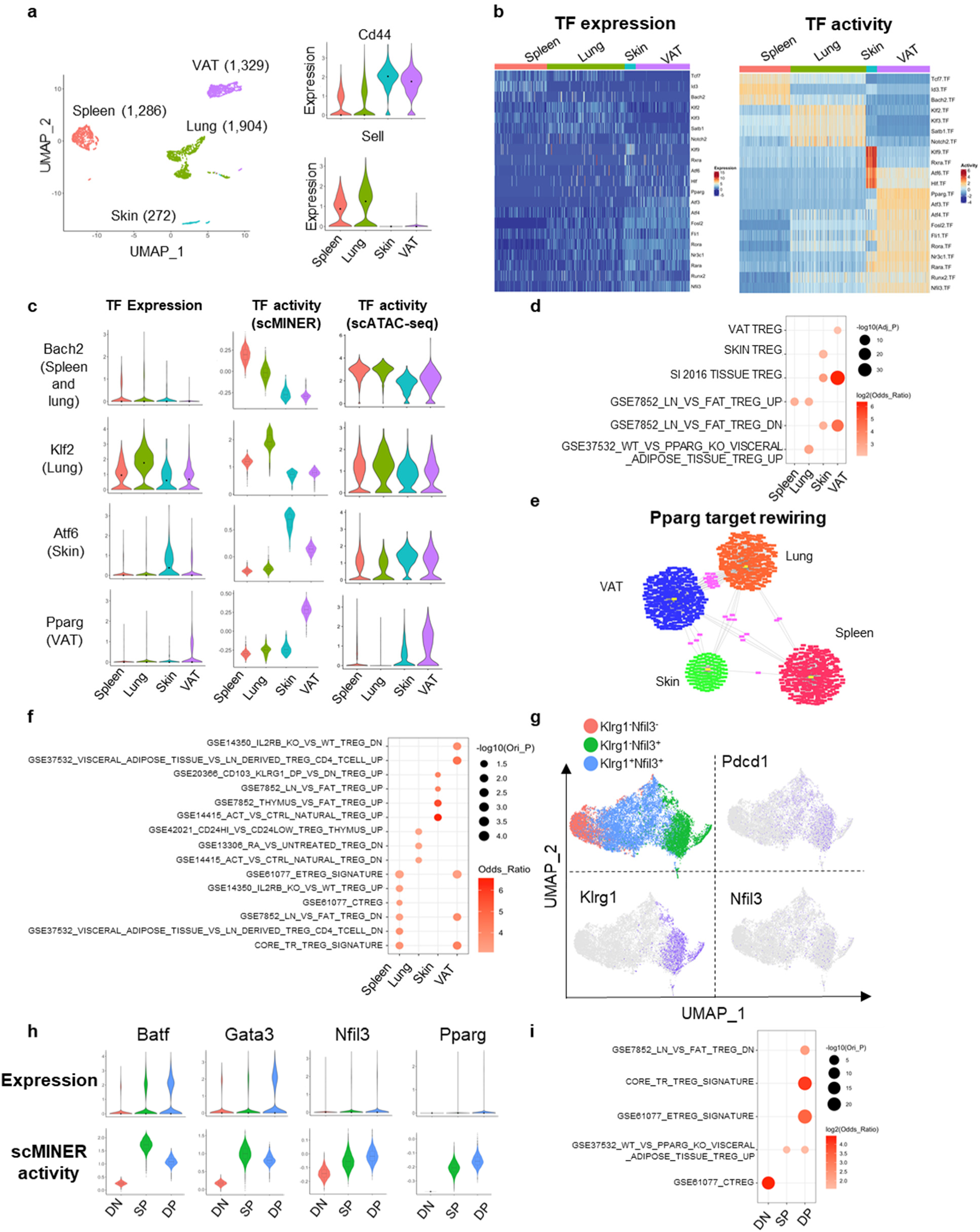
Tissue-specific differentiation of Treg identified by scMINER. **a,** MICA MDS clustering of mouse Foxp3^+^ regulatory CD4^+^ T cells (GSE130879) isolated from spleen, lung, skin and visceral adipose tissue (VAT). The expression of *Cd44* and *Sell* is visualized by violin plots. **b,** Heatmap visualization of predicted activity of top TFs in each cell from spleen, muscle, colon and VAT Treg cells. **c,** Violin plot visualization of *Bach2, Klf2, Atf6* and *Pparg* expression, scMINER activity, scATAC gene activity in spleen, muscle, colon and VAT Treg cells. scATAC gene activity was by signac R package based on GSE156112. **d,** Functional pathway enrichment of a union of top 50 TFs and top 200 SIGs in each tissue Treg cells based on t value from Student’s t test. **e,** The regulons of Pparg in spleen (red), lung (orange), skin (green) and VAT (blue) Treg cells. Regulons shared by 2 or more cell types are highlighted as pink. **f,** Functional pathway enrichment of Pparg regulons in spleen, muscle, colon and VAT Treg cells. **g,** MICA MDS clustering of mouse Klrg1^−^Nfil3^−^, Klrg1^−^Nfil3^+^, Klrg1^+^Nfil3^+^ Treg cells isolated from spleen (GSE130879). *Pdcd1*, *Klrg1* and *Nfil3* expression are visualized on UMAP. **h,** Violin plot visualization of Batf, Gata3, Nfil3 and Pparg expression and activity in spleen, muscle, colon and VAT Treg cells. **i**, Functional pathway enrichment of a union of top 50 TFs and top 200 SIGs in each stage of spleen Treg cells.

The prediction of selective scMINER activity, but not expression, of *Bach2* and *Pparg* in spleen and VAT Treg cells could be validated by another independent scRNA-seq study that profiled spleen, colon, muscle, and VAT Treg cells^61^ (**Supplementary Fig. 9c, d**). This further indicates the robustness and reproducibility of scMINER in identifying regulators of tissue-specific Treg cells from different studies. Moreover, the top scMINER-predicted TF and SIG drivers from skin and VAT Treg cells were significantly enriched in the core tissue resident Treg signature. They were enriched in the skin and VAT-specific signatures, respectively (**Fig. 6d**). These results indicated the top TF and SIG drivers predicted by scMINER could faithfully reveal tissue Treg differentiation reported in the literature.

To validate whether TF regulons predicted by scMINER also reflect TF transcriptional activity, we compared the scMINER-predicted regulons with footprint genes detected by ATAC-seq profiles of corresponding spleen and VAT Treg cells and indeed found a significant overlap of TFs between them (**Supplementary Fig. 9e**). *Pparg* is known as a master regulator of VAT Treg cells^64, 65^, and its transcription activity was correctly captured by scMINER in VAT (**Fig. 6c and Supplementary Fig. 9d**). Intriguingly, *Pparg* regulons are distinctly rewired in different tissues (**Fig. 6e**). The *Pparg* regulons in VAT Treg cells are significantly enriched in the VAT/Fat Treg signatures based on ATAC-seq analysis^63^ (**Fig. 6f**). Notably, predicted *Pparg* regulons in the spleen are also enriched in the core tissue Treg signature (**Fig. 6f**). This finding suggests *Pparg* may have an essential role in regulating early tissue Treg development and is consistent with previous studies^66, 67^. To further investigate this, we analyzed a scRNA-seq dataset of tissue Treg cell precursors (KLRG^−^NFIL3^+^ and KLRG^+^NFIL3^+^) in the spleen^66^ by scMINER (**Fig. 6g**). Tissue Treg cell precursors also express higher *Pdcd1*, *Klrg1* and *Nfil3* than KLRG^−^NFIL3^−^ cells. *Batf* is known to regulate the generation of tissue Treg cell precursors^66^. scMINER identified increased activity of *Batf*, *Gata3*, and *Nfil3* in the two tissue Treg cell precursors, consistent with the original study^66^(**Fig. 6h**). Indeed, *Pparg* activity was enhanced in these tissue Treg cell precursors despite its low expression, which supports that *Pparg* could regulate a common step of early tissue Treg cell generation in the spleen through its tuned regulons in the spleen, consistent with our observation above. Apart from *Pparg*, top TF and SIG drivers in the KLRG^+^NFIL3^+^ tissue Treg precursors are enriched in the core tissue resident Treg cell signatures and *Pparg* related gene expression (**Fig. 6i**), indicating that scMINER can reveal the drivers underlying the early generation of tissue Treg cell precursors in the spleen.

Similar to TF drivers, top SIG drivers for each tissue specific Treg cell are also highlighted (**Supplementary Fig. 9f**). Among these signaling drivers, *Ccr1* was demonstrated to possibly contribute to VAT Treg accumulation^65^, which was correctly captured by scMINER. In addition, a common epigenetic and transcriptional hallmark of tissue Treg cells is the upregulation of IL-33 receptor ST2 (*Il1rl1*)^68^. Treg cells from all three other Treg cells upregulated ST2 activity compared to the spleen. Still, VAT Treg cells have the highest ST2 activity (**Supplementary Fig. 9f**), which supports the critical role of the IL-33-ST2 axis in pan-tissue Treg cell generation. Moreover, other chemokine receptors such as *Ccr4* and *Ccr5*, which are essential in tissue Treg cell migration^68, 69^, have also been uncovered by scMINER. Together, these results indicate that scMINER can capture the upstream signals that modulate tissue Treg cell specification.

## Discussion

We have developed an integrative computational framework for unsupervised clustering and reconstruction of cell-type specific intracellular networks that enables the identification of hidden drivers directly from scRNA-seq data. Our scMINER framework takes advantage of mutual information for nonlinear measurements of intrinsic relationships among cells and genes. For clustering, scMINER also leverages an ensemble dimension reduction approach based on a reasonable assumption of the nature of biological dynamics: that a wider range of dynamics should be observed from a large population of cells of the same cell type. Using benchmarking datasets with ground-truth labels, we demonstrated that scMINER outperforms the state-of-the-art methods in single-cell clustering.

From scRNA-seq data alone, scMINER is able to uncover hidden TF and SIG drivers underlying cell states, which have not been captured by differential gene expression analysis but have been experimentally validated as key drivers. Further, the scMINER-inferred TF regulons are significantly overlapped with the targets defined from ATAC-seq footprinting of the same cell type, suggesting high accuracy of the scMINER-derived TF networks. For TF driver prediction, we benchmark scMINER with SCENIC and demonstrate that scMINER identifies known TF drivers that SCENIC fails to reveal due to the lack of TF binding motif information or low expression of TFs. Importantly, scMINER is the only method that can predict SIG drivers from scRNA-seq data. Together, scMINER provides a new toolbox to dissect the TF regulatory and signaling networks and pinpoint hub drivers underlying cell lineage differentiation and specification from single-cell omics data.

In addition to improving clustering accuracy, scMINER provides insights that could only be gleaned using an activity analysis pipeline. In particular, the established lineage markers of some subsets of lymphocytes, especially regulatory T cells, were missed in the conventional expression-based analysis, due to significant gene dropouts. In contrast, activity-based scMINER analyses rescue the dropout and reveal the hidden drivers and rewiring of their regulons in each cell state. Our results show that the importance of these cell-state drivers is consistent with their role reported in literature and these top drivers can be faithfully recapitulated in multiple scRNA-seq datasets profiled in similar experimental contexts. Moreover, scMINER provides quantitative activity assessment for > 6,000 TF and SIG proteins in a single experiment; it outperforms traditional low-throughput methods examining protein expression and activity, such as flow cytometry; and it overcomes the limitations of TF motif based SCENIC activity inference. Thus, activity-based scMINER analyses provide higher accuracy, reproducibility and scalability for inferring cellular transcriptional networks in each cell state from scRNA-seq data.

Importantly, the reasons why activity-based scMINER analyses outperforms single cell expression-based analysis in the identification of multiple known transcription factors and signaling pathways go beyond the rescue of dropout. For instance, in regulating CD8^+^ T cell exhaustion and Treg cell tissue specification, *Batf* is an example of hidden driver because its expression is largely comparable among three subsets of CD8^+^ exhausted T cell subsets. However, its role acting in opposing Tpex generation is reported in the literatures^33, 45^. Thus, the identification of *Batf* has a more important role for effector-like TCF1^−^TIM3^+^ cells by scMINER, which stripped the disguise of its expression profile. The rewiring of the TF targets shown in different subsets of CD8^+^ T or Treg cells also suggests the necessity of studying cell-cluster specific TF regulons rather than the invariant TF regulons predicted by SCENIC, which could help in understanding unique transcriptional regulation in novel cell states and the molecular mechanism underlying the ‘hidden’ drivers. Since CD8^+^ T cell exhaustion and Treg cell development in tissues are both critical in regulating tumor and infection progression, targeting these top master regulators could help relieve these diseases and open the door for new combinational immunotherapies using current checkpoint blockade strategies.

While scMINER is a robust and powerful tool for single-cell clustering and network analysis, it has several limitations. First, the nonlinear MI estimation of cell-cell or gene-gene similarities is time-consuming, especially with a large number of cells. For clustering, scMINER doesn’t select top variable genes to retain information that can separate close cell states. Computing platforms supporting parallelism may help improve the efficiency. For network inference of big clusters, downsampling is one solution, but a MetaCell-type approach is preferred because it also helps improve the gene coverage by aggregating expression in multiple cells. Second, although scMINER provides silhouette analysis to guide selection of the optimal number of clusters, determining the number of clusters is still quite challenging, thus a manual and problem-dependent task.

In summary, we report the development and application of a novel scRNA-seq analytic pipeline, which utilizes mutual information and network-based inference to complement gene expression for better clustering and predicting the importance of TF and SIG drivers in each cell type. The MICA clustering algorithm in scMINER can also be applied to other high-dimensional data, including scATAC-seq, bulk transcriptomics and proteomics, and spatial omics. While our examples focused on the immune system, the scMINER tool could be effectively applied to any other systems profiled by scRNA-seq.

## Methods

### Data compilation and preprocessing

We summarize the 11 single-cell data sets used for accessing the clustering methods in Table 1. We filtered out genes detected in less than three cells, and cells that express less than 200 genes. The gold (Yan, Goolam, Pollen, Kolod) and silver standard data sets (Usoskin, Klein, Zeisel) were normalized using the same methods reported in SC3^11^. In addition, the Buettner data set was normalized using FPKM; Chung data set was normalized with TPM; all the UMI-based data sets were normalized to 10,000 reads per cell. We then performed a natural log transformation for all the normalized data. In addition, we extracted highly variable genes using the default parameters for the clustering analysis by Seurat and Scanpy as recommended in the tutorials for PCA dimension reduction. We only used the known cell labels afterward to access the clustering results.

The dataset used in Fig. 3 was based on PBMC dataset generated by Zheng et al.^15^. It has 20,000 PBMCs purified via well-known cell surface markers with each subpopulation of 2,000 cells. To create a more purified simulation dataset, CD4+ T helpers, total CD8+ cytotoxic T, and CD34+ cells (HSPCs) were removed with the remaining 14,000 cells forming seven mutually exclusive cell types.

### Mutual information estimation

Cellc using MI:

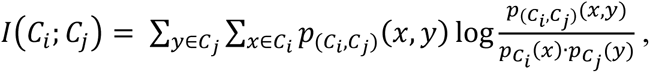

where *C_i_* and *C_j_* are the *i*-th and *j*-th rows in matrix *M*. Since normalized gene expression values are continuous, we use a binning approach^70^ for discretizing the expression values for the joint and marginal probability calculations, where the bin size *b* is defined as 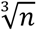, n being the total number of genes. We then define a normalized distance between cells as

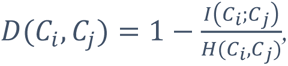

where *H*(*C_i_*, *C_j_)* is the joint entropy of cells *C_i_* and *C_j_*.

### Ensemble dimension reduction

For data sets with more than t (default is 5,000) cells (according to the clustering performance evaluation), we first represent MI matrix *M* as a *k*-nearest neighbor graph *G_l_* and then adopted node2vec^71^ algorithm to embed *G_l_* to a *d* dimensional space, where *k* and *d* are predefined parameters to control the scale of the local structure to explore and the number of low dimensional components to transform, respectively. The embedding is performed for a range of *d* from 8 to 96 with step size four followed by consensus clustering to reduce the randomness caused by selecting a fixed *d* in a single clustering run. We use classical multidimensional scaling (MDS) to preserve the intercell distances for data sets with less than or equal t cells. We set the number of components to be a fixed number of 19, as explained in the Main section (**Fig. 2d**). As MDS requires storing the entire MI matrix in the memory, to enable MI estimation for large datasets, we parallelized the calculation in four steps: 1) partitioning *M* into *M_i,j_* of a fixed number of cells, 2) MI estimation for every pair of *M_i,j_* in parallel, 3) merging MI matrices, and 4) normalization using equation (1).

### Cell clustering

To ensure reasonable running time, we build a k-nearest neighbor (kNN) graph *G_2_* again after embedding *G_l_* to a *d* dimensional space and use the Louvain algorithm as the default clustering method. Instead of using mutual information as the distance function, we use Euclidean distance to determine the similarity of cells on the *d* dimensional space. Euclidean distance has much fewer arithmetic operations than MI; thus, we adopt the KD-tree algorithm^70^ to construct an exact kNN graph *G_2_* instead of an approximate graph *G_l_*. Similar to SC3, we use k-means clustering as the default method on the MDS transformed space. However, K-means clustering is sensitive to the centroid seed initialization and produces very different results for each run. To reduce randomness, we run k-means clustering ten times by default and use consensus clustering to aggregate multiple runs of k-means clustering results.

### Determination of the optimal number of clusters

We compute a silhouette coefficient^17^ average over all the cells for the clustering results and rank the results by the coefficients. scMINER users may evaluate the top candidates to determine the optimal number of clusters with a priori knowledge of the biological context of the data set.

### Determination of parameters for graph-embedding in MICA

With the increasing complexity and depth of cellular neighborhood in the cell-cell distance space for large datasets, we speculate that the ability to explore the diverse local neighborhoods in a non-linear fashion is important to the clustering performance. Therefore, we adopt node2vec^71^, a non-linear graph-based dimension reduction approach with the flexibility in exploring diverse neighborhoods, to maximize the likelihood of preserving local neighborhoods of cells. Most of the current dimension reduction techniques rely on eigendecomposition of the appropriate data matrix, which unsatisfies the scalability requirement for large datasets. A feasible solution is to explore and preserve the local neighborhood of cells approximated by a graph representation with each node representing a cell. node2vec’s random walk-based approach with flexible parameters, which allows for unsupervised exploration of the local neighbors in a graph and offers scalability for large datasets without sacrificing much clustering performance. We choose a subset of parameters critical to the dimension reduction performance and perform the grid searches over a predefined range for each parameter independently using the Zheng dataset. The default parameters are selected by considering the clustering performance in terms of ARI, the elapsed wall time and the highest memory consumption.

### Performance analysis and parameter tuning

We compute the running time for each step of MICA for PBMC and Human Motor Cortex datasets using 25 CPU cores from a redhat linux machine (**Supplementary Fig. 3a**). The 1^st^ step (MI-kNN) takes more than half of the total running time due to the number of arithmetic operations of mutual information calculation for each cell pair. We also perform parameter tunings by grid searching in a predefined range of selected parameters for PBMC clustering analysis (**Supplementary Fig. 3b**). The parameters in MI-kNN and node2vec steps have the greatest influence on the clustering performance and running time. We chose the default parameters to balance the clustering performance, running time and memory usage.

### Overview of MINIE

MINIE allows users to reconstruct cell-type-specific GRNs for driver activity inference and target network rewiring analysis. MINIE takes inputs of a gene expression profile and cell cluster labels. It first filters the genes with all zero expressions on the cell cluster basis. Then MINIE invokes SJARACNe to reconstruct cell-type-specific transcriptional factor and signaling networks. While users can provide a list of known drivers (TFs or SIGs) as input, well-curated lists of drivers are included in the MINIE package for users’ convenience. To resolve the sparseness of the networks due to the scRNA-seq dropout effect, we fine-tuned SJARACNE parameters (e.g., two p-value thresholds) to consider a series of network properties, including the median number of targets, a power law-like degree distribution, etc. Finally, with the predicted targets of a driver for a cell cluster, MINIE calculates the driver activity by performing a column-wise normalization to ensure each cell is on a similar expression level, followed by averaging the expression of the driver’s target genes.

### Bulk ATAC-seq data analysis

ATAC-seq analysis was performed as described previously. Briefly, two × 50-bp paired-end reads we obtained from public datasets were trimmed for Nextera adaptor by trimmomatic (v0.36, paired-end mode, with parameter LEADING:10 TRAILING:10 SLIDINGWINDOW:4:18 MINLEN:25) and aligned to mouse genome mm10 downloaded from gencode release M10 (https://www.gencodegenes.org/mouse/release_M10.html) by BWA (version 0.7.16, default parameters). Duplicated reads were then marked with Picard (v2.9.4), and only non-duplicated proper paired reads were kept by samtools (parameter ‘-q 1 -F 1804’ v1.9). After adjustment of Tn5 shift (reads were offset by +4 bp for the sense strand and −5 bp for the antisense strand), we separated reads into nucleosome-free, mononucleosome, dinucleosome, and trinucleosome as previously described by fragment size and generated ‘.bigwig’ files by using the centre 80-bp of fragments and scaled to 30 × 10^6^ nucleosome-free reads. We observed reasonable nucleosome-free peaks and a pattern of mono-, di- and tri-nucleosomes on IGV (v2.4.13). All samples in this study had approximately 1 × 10^7^ nucleosome-free reads, indicating good data quality. Next, peaks were called on nucleosome-free reads by MACS2 (v2.1.1.20160309, with default parameters with ‘–extsize 200–nomodel’). To assure reproducibility, we first finalized nucleosome-free regions for each sample and retained a peak only if it called with a higher cut-off (MACS2 -q 0.05). We further generated consensus peaks for each group by keeping peaks presenting in at least 50% of the replicates and discarding the remaining, non-reproducible peaks. The reproducible peaks were further merged between samples if they overlapped by 100-bp; we counted nucleosome-free reads from each sample by bedtools (v.2.25.0). To identify the differentially accessible open chromatin regions (OCRs), we first normalized raw nucleosome-free read counts per million (CPM) followed by differential accessibility analysis by implementing the negative binomial model in the DESeq2 R package^72^. FDR-corrected *P-value* < 0.05, |log_2_ FC| > 0.5 were used as cut-offs for more- or less-accessible regions in TOX KO samples compared to their WT samples. Principal component analysis was performed using function prcomp in R. We then assigned the differentially accessible OCRs in the ATAC-seq data for the nearest genes to generate a list of DA genes using HOMER. This analysis identified 25,646 open chromatin regions (OCRs) with differential expression in TOX KO versus control cells (FDR < 0.05; |log_2_ FC (TOX KO/WT)| > 0.5).

### Motif analysis and footprinting of transcription factor binding sites

For motif analysis, we further selected 1,000 unchanged regions log2 FC < 0.5 and FDR-corrected P-value > 0.5 as control regions. FIMO from MEME suite (v4.11.3, ‘–thresh 1e-4– motif-pseudo 0.0001’)^73^ was used for scanning motif (TRANSFAC database release 2019, only included Vertebrata and not 3D structure-based) matches in the nucleosome-free regions, and two-tailed Fisher’s exact test was used to determine whether a motif was significantly enriched in differentially accessible compared to the control regions. To perform footprinting analysis of transcription factor binding site, the RGT HINT application was used to infer transcription factor activity and plot the results^51^.

### Single-cell ATAC-seq processing and data analysis

All preprocessing steps were performed using “Cell Ranger ATAC version 1.2.0” (10X Genomics). Read filtering, alignment, peak calling, and count matrix generation from fastq files were done per sample using ‘cellranger-atac count’. Reference genome assemblies mm10 (refdata-cellranger-atac-mm10-1.2.0) provided by 10xGenomics were used for samples. All further analysis steps were performed in R (Version 4.0.0). Fragments were loaded into R using the package Seurat^10^. The R package Signac (version 1.3.0, https://github.com/timoast/signac) was used for normalization and dimensionality reduction. The peak-barcode matrix was then binarized and normalized using the implementation of the TF-IDF transformation described in (RunTFIDF (method = 1)). Subsequently, singular value decomposition was run (RunSVD) on the upper quartile of accessible peaks (FindTopFeatures (min.cutoff = ‘q75’)). The first 20 components from the SVD reduction were used for secondary dimensionality reduction with UMAP.

### Gene activity scores of scATAC-seq data

To calculate gene activity scores, gene body coordinates were first obtained by using the command genes (TxDb.Mmusculus.UCSC.mm10.knownGene) from the package GenomicFeatures in R. The coordinates were filtered for normal chromosomes (keepStandardChromosomes (pruning.mode = ‘coarse’)) and extended by 2,000 bp upstream of the transcription start sites to include promoter regions (Extend(upstream = 2000)). Then, the command ‘FeatureMatrix’ from the Signac package was used with the ‘features’ parameter set to the extended gene coordinates, to sum up the number of unique reads within gene regions for each cell. The above steps can be performed using the wrapper function GeneActivity. Eventually, these gene activity scores were log-normalized and multiplied by the median read counts per cell (nCount_Reads) with the command NormalizeData(normalization.method =’ LogNormalize’,scale.factor = median(nCount_Reads). Normalized gene activities were capped at the 95th quantile for plotting.

### Estimation of transcription factor activity with chromVAR

Transcription factor (TF) activities for each cell were measured using chromVAR (Schep et al., 2017). TF position weight matrices were downloaded from the Homer website (http://homer.ucsd.edu/homer/custom.motifs). Signac was used to build a motif-peak matrix for all peaks in the murine and human datasets (CreateMotifMatrix) using reference genomes from the packages BSgenome.Mmusculus.UCSC.mm10 and BSgenome.Hsapiens.UCSC.hg19, respectively. After assembling and adding the Motif object to the Seurat object (CreateMotifObject, AddMotifObject), information on the base composition was calculated for each peak (RegionStats). Eventually, the wrapper function ‘RunChromVAR’ was called to obtain chromVAR deviation z-scores. ChromVAR deviation z-scores below the 5th and above the 95th quantile were capped for plotting.

### SCENIC regulon analysis

Previously published scRNA-seq data of CD8^+^ T cells from LCMV infection model (GSE122712, GSE119940), tissue-specific Treg cells (GSE130879) were used for SCENIC analysis^6^, with raw count matrix as input. Briefly, the co-expression network was calculated by GRNBoost2, and RcisTarget identified the regulons. Next, the regulon activity for each cell was scored by AUCell. For some regulons, AUCell thresholds were manually adjusted as recommended by the SCENIC developers. Finally, the activity of each transcription factor was visualized by heatmap among all the cell clusters.

### Calculate activity from MetaCell

Raw count matrix was imported by mcell_import_scmat_tsv function in metacell R package and metacell membership was calculated by its default pipeline. Cells with Log2 (library normalized) gene expression was assigned the metacell membership and average gene expression of cells with the same membership was calculated to form a gene x metacell pseudobulk matrix. Based on the TF and SIG list in SJARACNE, the gene x metacell matrix was exported by generateSJARACNeInput function to for generating TF and SIG network by SJARACNE. The activity of each cell was further calculated with cal.Activity function with es.method=’weightedmean’.

### Software packages

For comparing the clustering performance with Seurat, SC3, and Scanpy, we used the following packages: (i) Seurat version 4.0.3 from CRAN (https://cran.r-project.org/web/packages/Seurat/index.html); (ii) SC3 version 1.18.0 from Bioconductor (https://bioconductor.org/packages/release/bioc/html/SC3.html); (iii) Scanpy version 1.6.0 from GitHub (https://github.com/theislab/scanpy); for approximated nearest neighbor graph construction, we used PyNNDescent version 0.5.2 (https://github.com/lmcinnes/pynndescent), a Python implementation of NNDecent algorithm^74^; for graph embedding, we used PecanPy^75^, an efficient Python implementation of node2vec^71^; we used an open-source compiler Numba (http://numba.pydata.org/) for translation of our Python and NumPy implementation of mutual information calculation into fast machine code.

## Data availability

All the data sets in **Supplementary Tables 1 and 2** were downloaded from the accession numbers provided in the original publication.

## Code availability

The source code for scMINER is available online at https://github.com/jyyulab/scMINER. The documentation with a tutorial is available online at https://jyyulab.github.io/scMINER.

## Acknowledgements

We thank the members of the Yu Lab for testing and improving scMINER and Keith A. Laycock for scientific editing. This work was supported in part by National Institutes of Health grants R01GM134382 (to J.Y.), U01CA264610 (to J.Y.) and R35CA253188 (to H.C.), and by the American Lebanese Syrian Associated Charities. The content is solely the responsibility of the authors and does not necessarily represent the official views of the National Institutes of Health.

## Author contributions

D.L., H.S., and J.Y. conceived the project. D.L. and H.S. designed the computational method, wrote software packages, and carried out analyses. C.Q. contributed to analyses and software development. C.B., J.P.V., A.K., M.R., and M.B. contributed to software development. Q.P., Y.D., Z.X., I.R., X.Y., X.H., and L.Y. assisted with analysis and software testing. K.K.Y. provided computational insights. H.C. provided biological insights. D.L., H.S., C.Q., and J.Y. wrote the manuscript. J.Y. supervised the project.

## Supplementary Information

**Supplementary Figure 1.**
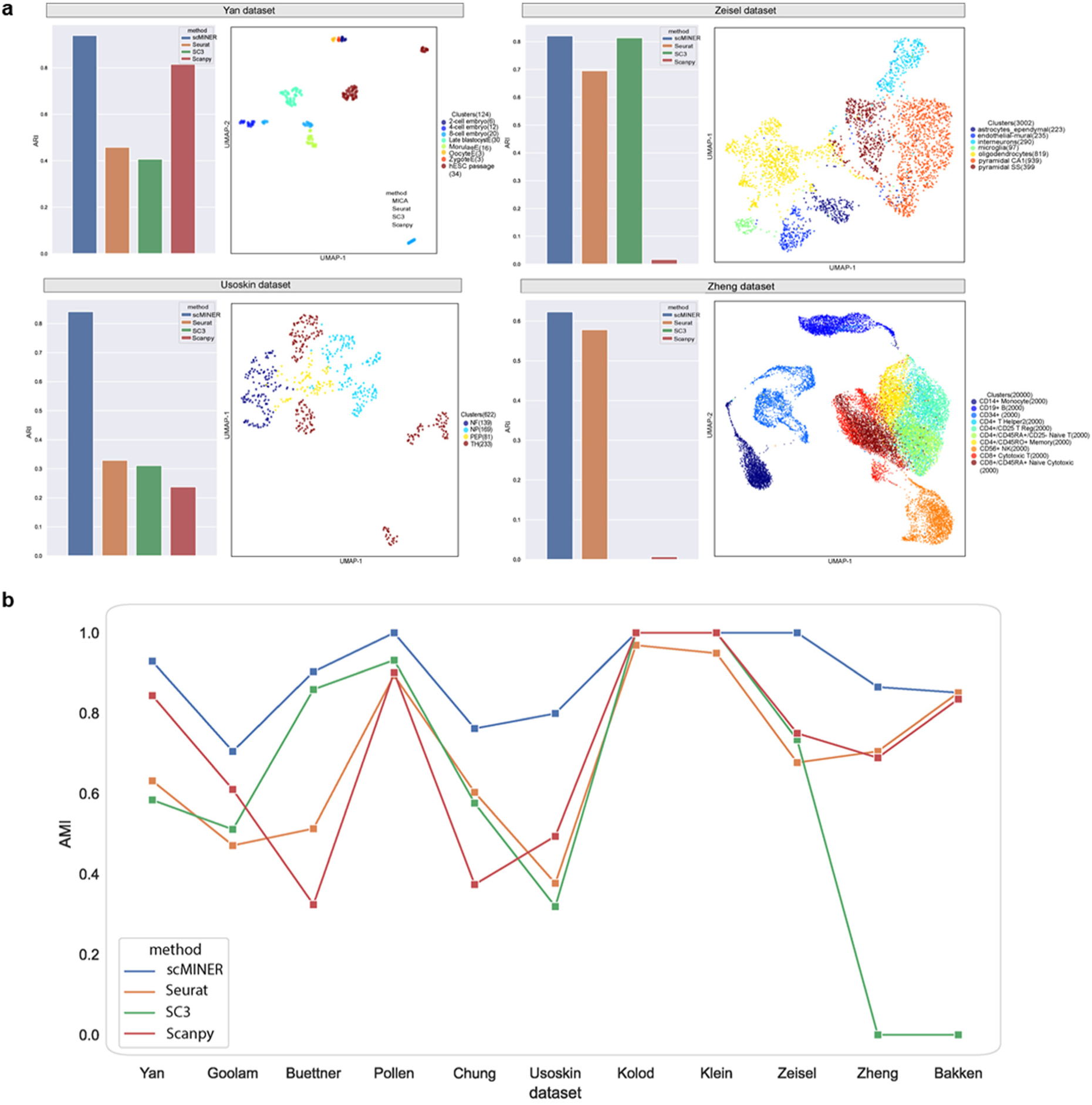
scMINER clustering performance evaluation using AMI and true label projection on four datasets. **a,** ARI bar plots and UMAP plots of scMINER clustering results annotated using true labels on Yan, Zeisel, Usoskon, and Zheng datasets. **b,** Clustering performance of scMINER, Seurat, SC3 and Scanpy measured by adjusted mutual information (AMI).

**Supplementary Figure 2.**
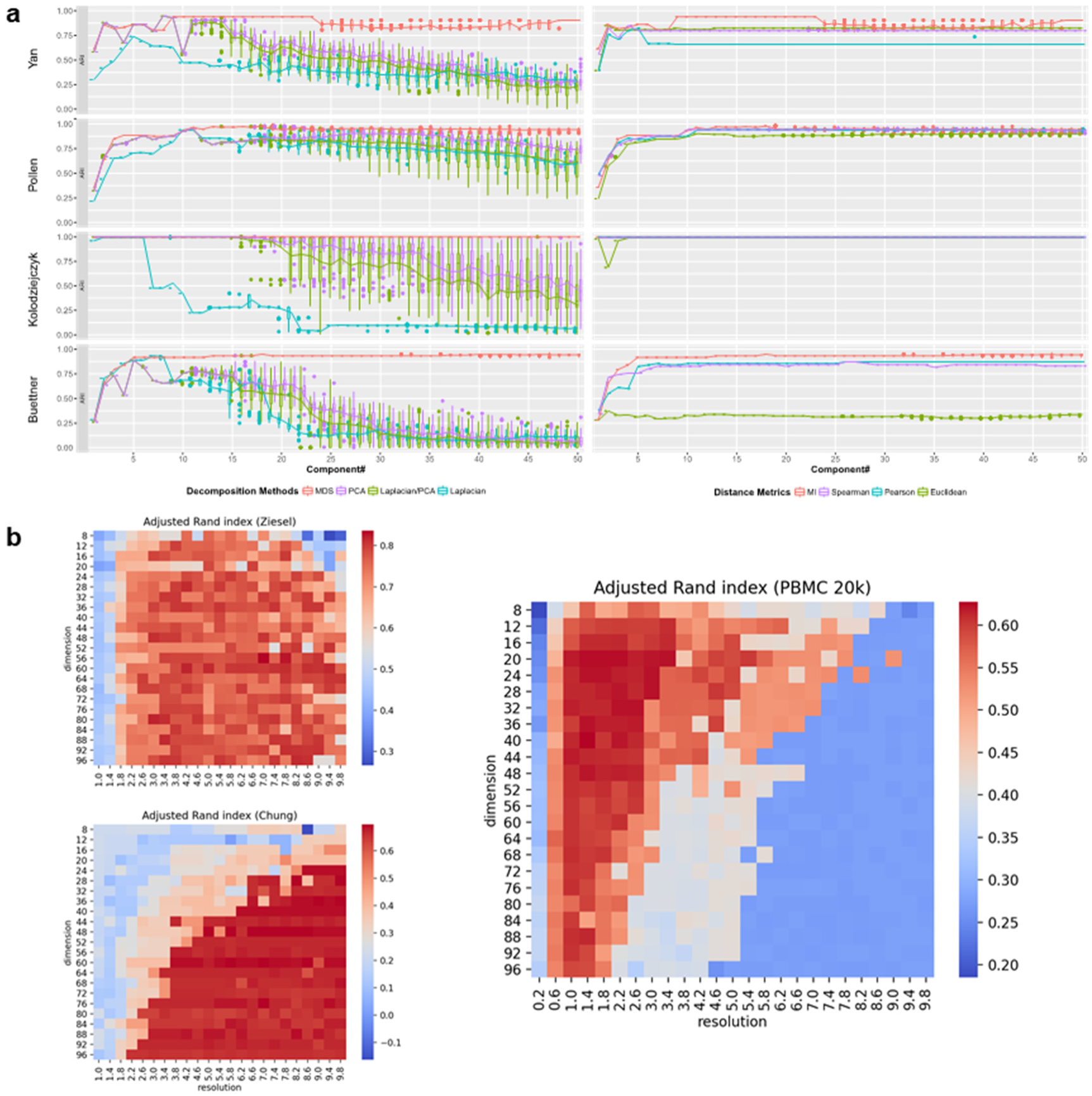
Effect of distance metrics and parameters on the clustering performance. **a,** Clustering performance comparison using four distance metrics (left) and four dimension reduction methods (right) on Yan, Pollen, Kolodziejczyk, and Buettner datasets. **b,** Clustering performance in term of ARI with respect to dimension and resolution parameters.

**Supplementary Figure 3.**
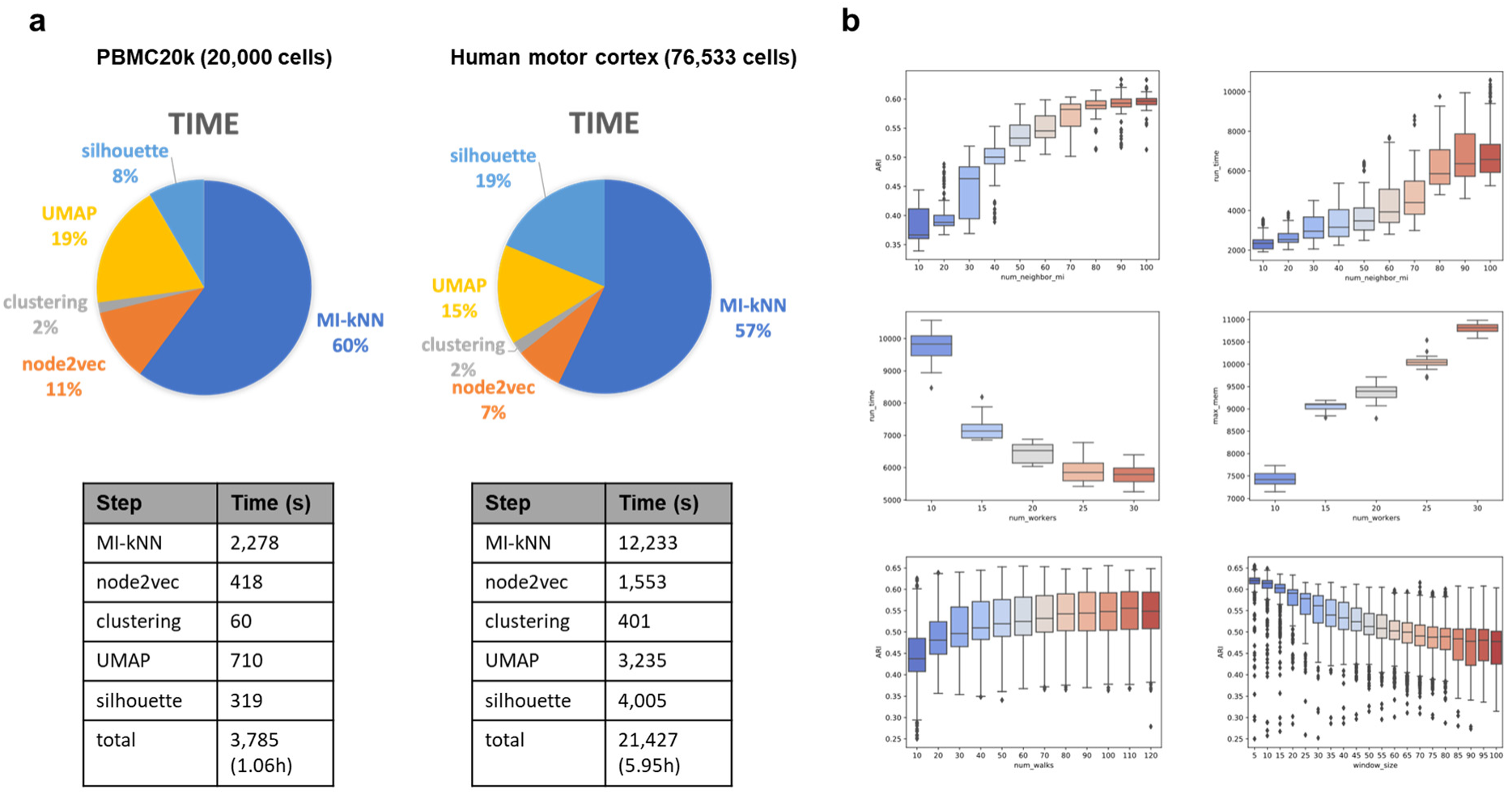
MICA computing resource usage analysis for PBMC (Zheng) and Human Motor Cortex (Bakken) datasets. **a,** Run time for each step of MICA for PBMC20k and human motor cortex datasets using 25 cores. **b,** ARI, run time and memory consumption for PBMC with respect to some important parameters, e.g., number of workers, number of neighbors in building MI-kNN, and node2vec window size, etc.

**Supplementary Figure 4.**
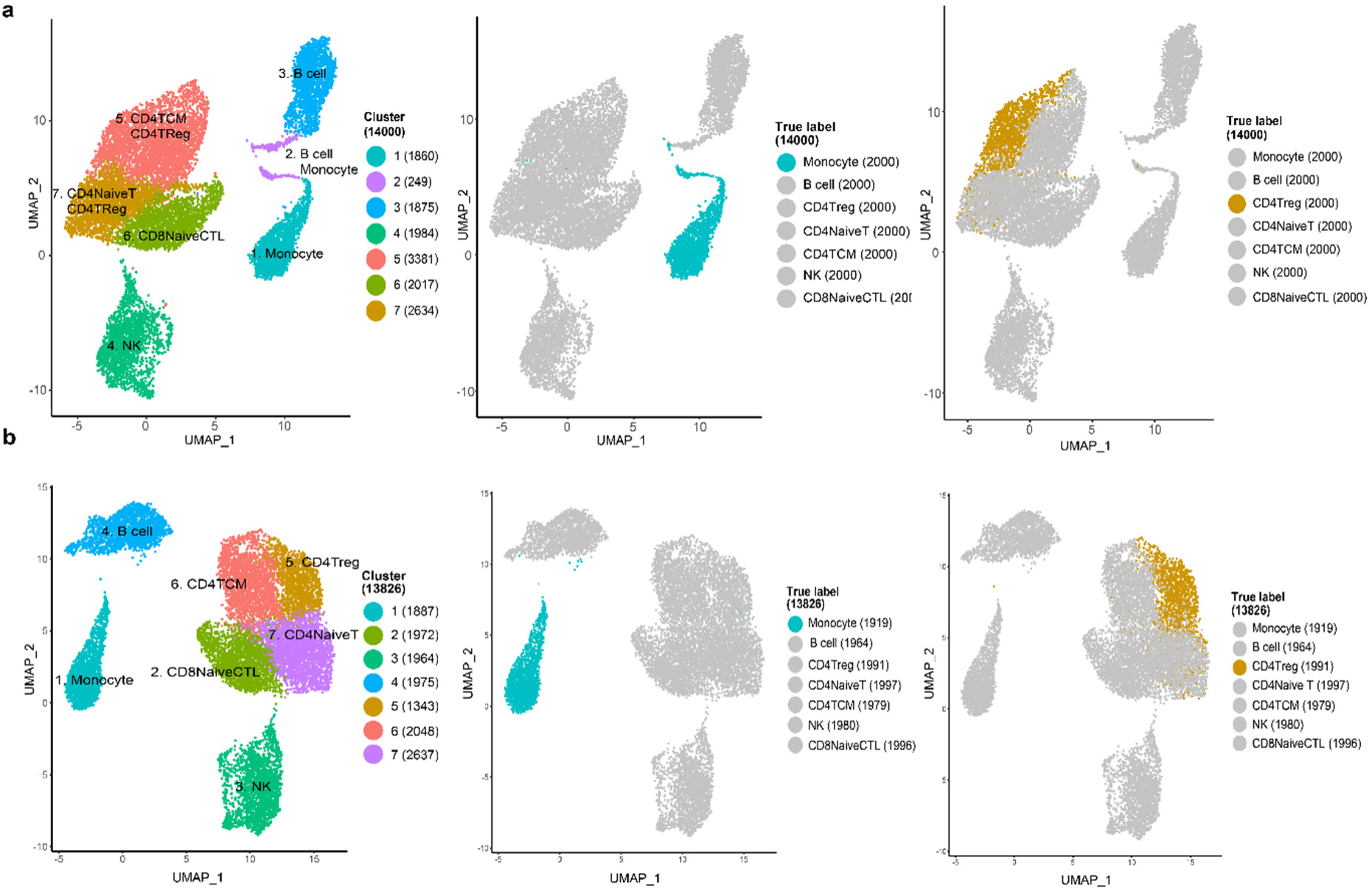
Effect of CP10K and CPM normalization on the clustering result of Zheng dataset. **a,** UMAP plots of all 7 clusters using count per 10K (CP10K) for normalization. **b,** UMAP plots of all 7 clusters using count per million (CPM) for normalization.

**Supplementary Figure 5.**
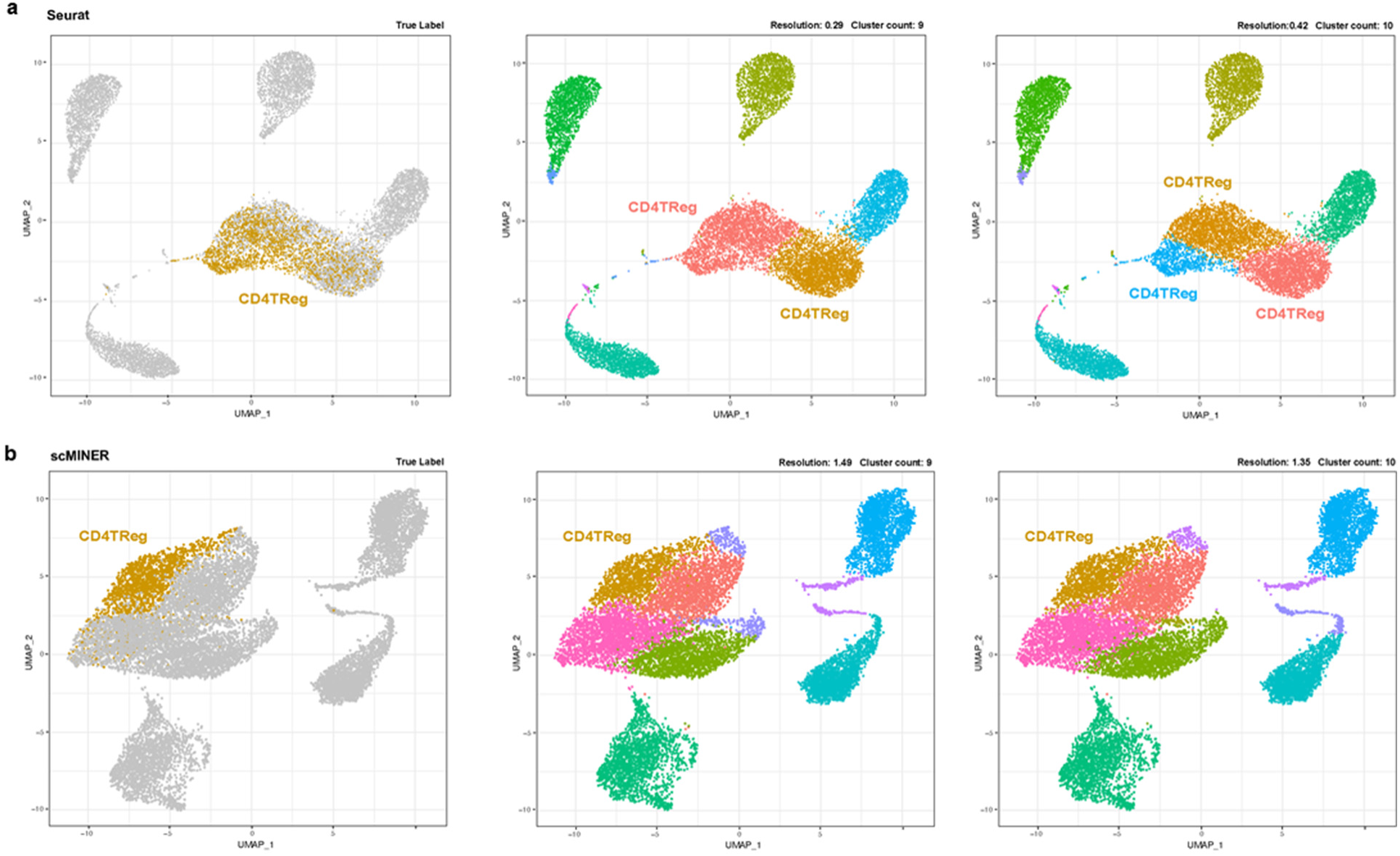
Comparison of scMINER and Seurat CD4Treg cell distribution on UMAPs with respect to the changing of clustering resolution. **a,** CD4Treg cell distribution on Seurat clusters with respect to the increasing number of resolution and cluster count. **b,** CD4Treg cell distribution on scMINER clusters with respect to the increasing number of resolution and cluster count.

**Supplementary Figure 6.**
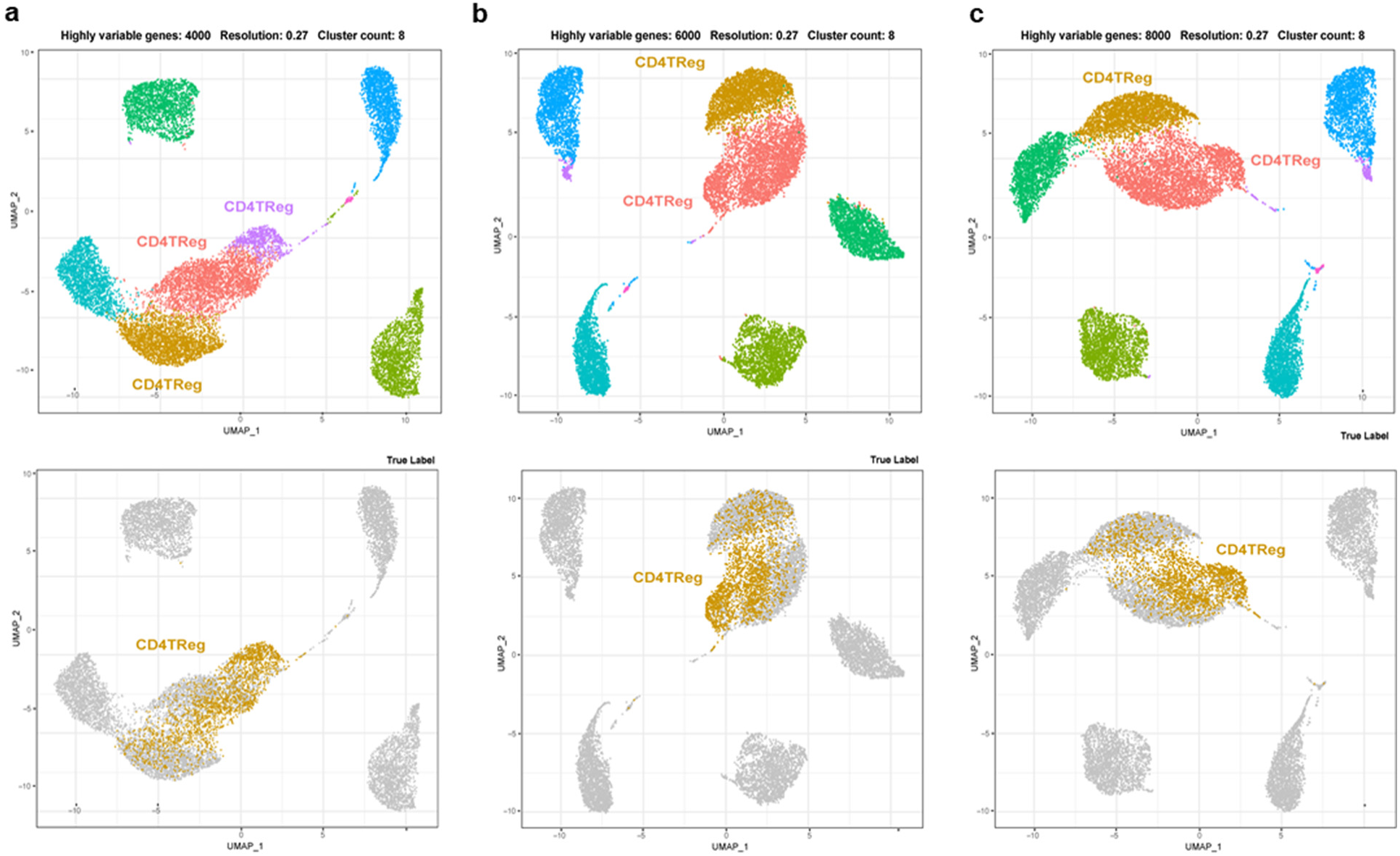
CD4Treg cell distribution on Seurat clusters with respect to the changing of the number of highly variable genes. a-c, CD4Treg cells are distributed in three Seurat clusters with 4,000 (a), 6,000 (b) and 8,000 (c) highly variable genes.

**Supplementary Figure 7.**
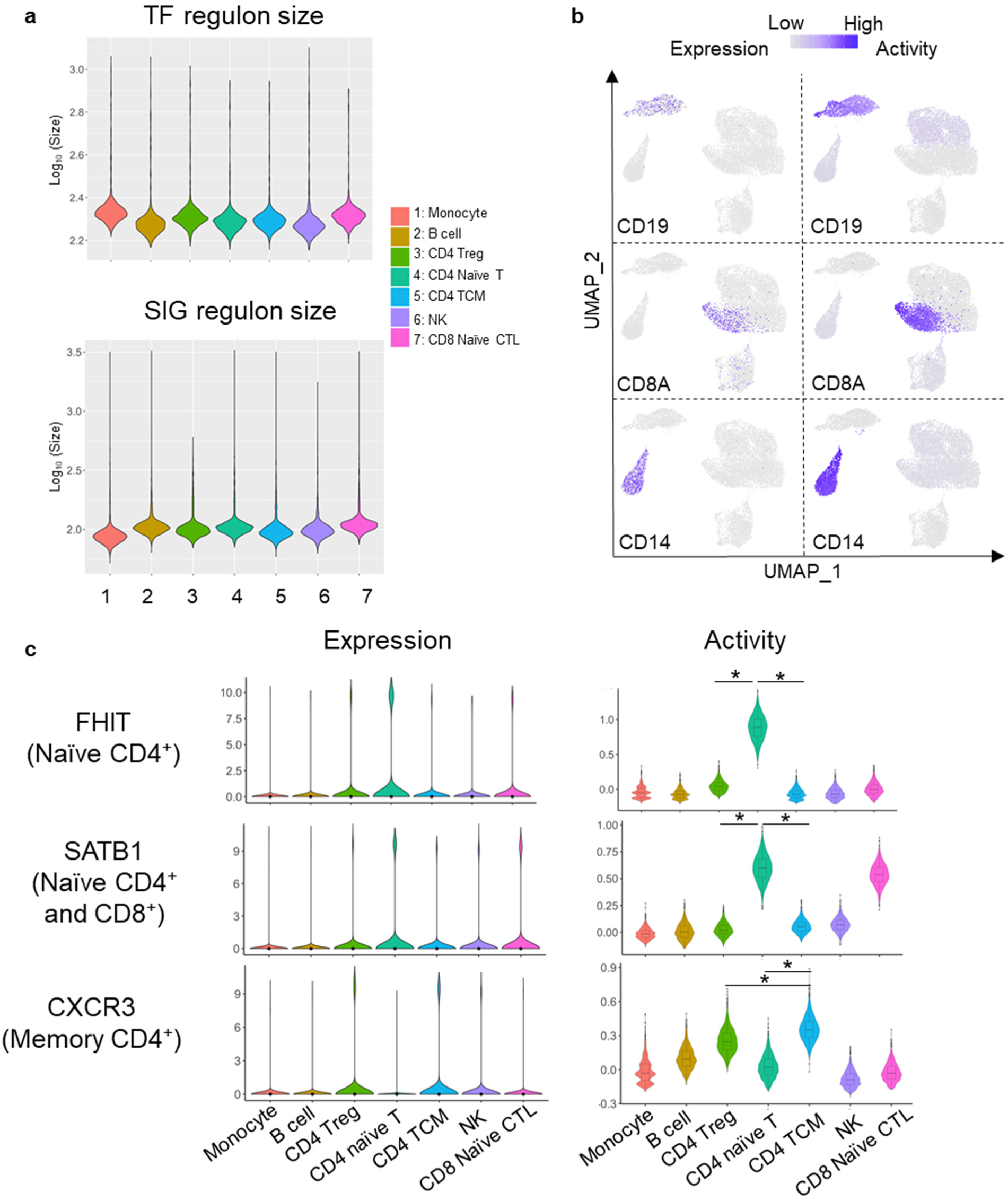
scMINER measures the activity of cell type-specific markers in PBMC. **a,** Mean TF and SIG regulon sizes in 7 sorted cell populations from PBMC scRNA-seq data. **b,** Expression and activity of CD19, CD8A and CD14 on UMAP using PBMC scRNA-seq data. **c,** Violin plot visualization of FHIT, SATB1 and CXCR3 expression and scMINER activity in 7 sorted cell populations from PBMC scRNA-seq data. *, *P* < 2e-16.

**Supplementary Figure 8.**
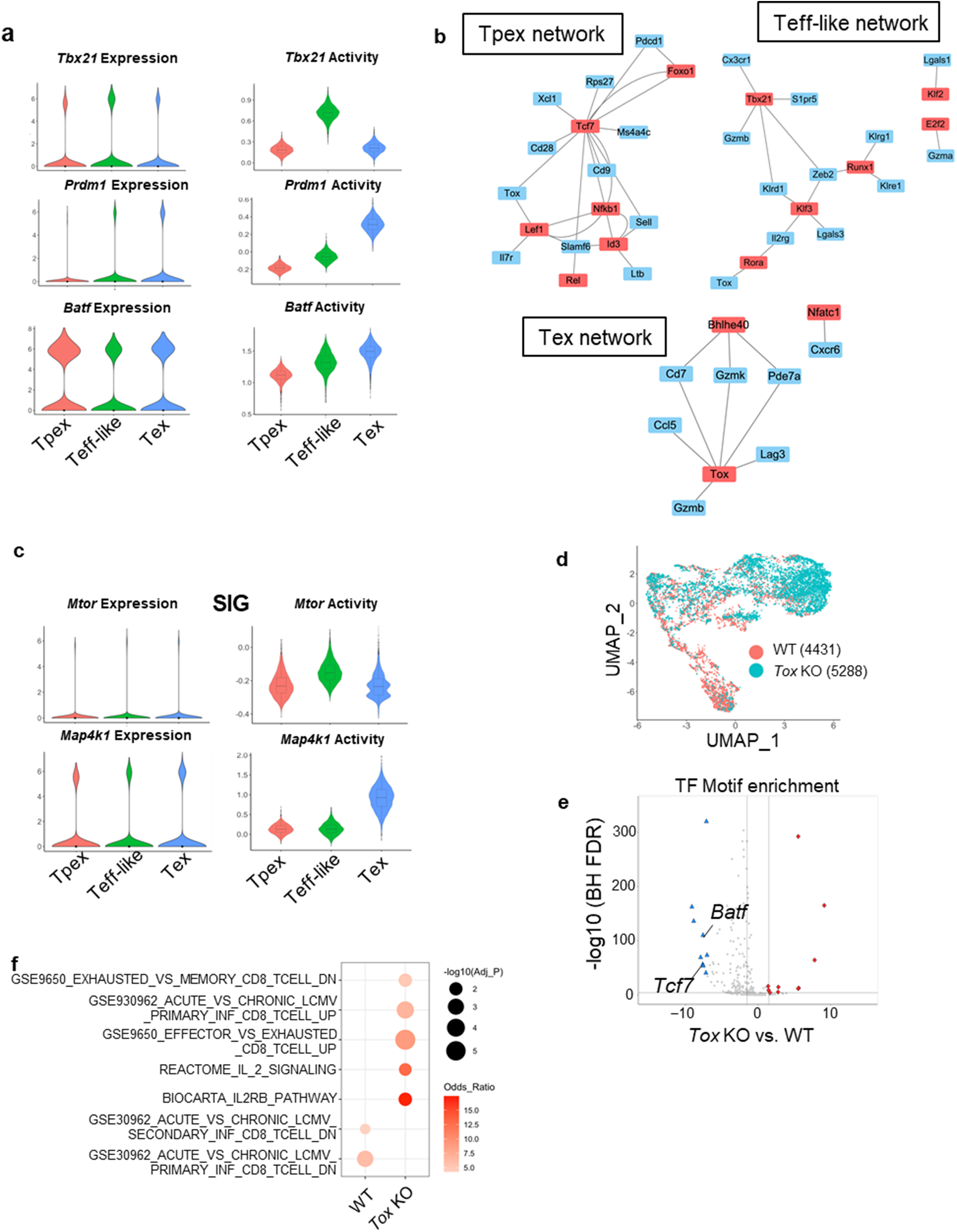
scMINER reveals drivers in wild-type and gene-perturbed CD8^+^ T cells during chronic infection. **a,** Violin plot visualization of Tbx21, Blimp1 and Batf expression and activity in 3 subsets of CD8^+^ T cells. **b,** GRNs for Tpex cells, Teff-like cells and Tex cells. Key TFs shown in Fig. 5b for each CD8^+^ T cell subset are highlighted in red. **c,** Violin plot visualization of *Mtor* and *Map4k1* expression and activity in 3 subsets of CD8^+^ T cells. **d,** UMAP visualization of wild type and *Tox* deficient CD8^+^ T cells in chronic infection (GSE119940). The numbers in the bracket indicates the cell numbers of each genotype. **e,** TF motif enrichment analysis for *Tox* deficient vs. wild-type CD8^+^ T cells using an ATAC-seq dataset (GSE132986). BH FDR, the Benjamini-Hochberg false discovery rate. **f,** Functional pathway enrichment of a union of top 50 TFs and top 200 SIGs predicted by scMINER for wild type and *Tox* deficient CD8^+^ T cells.

**Supplementary Figure 9.**
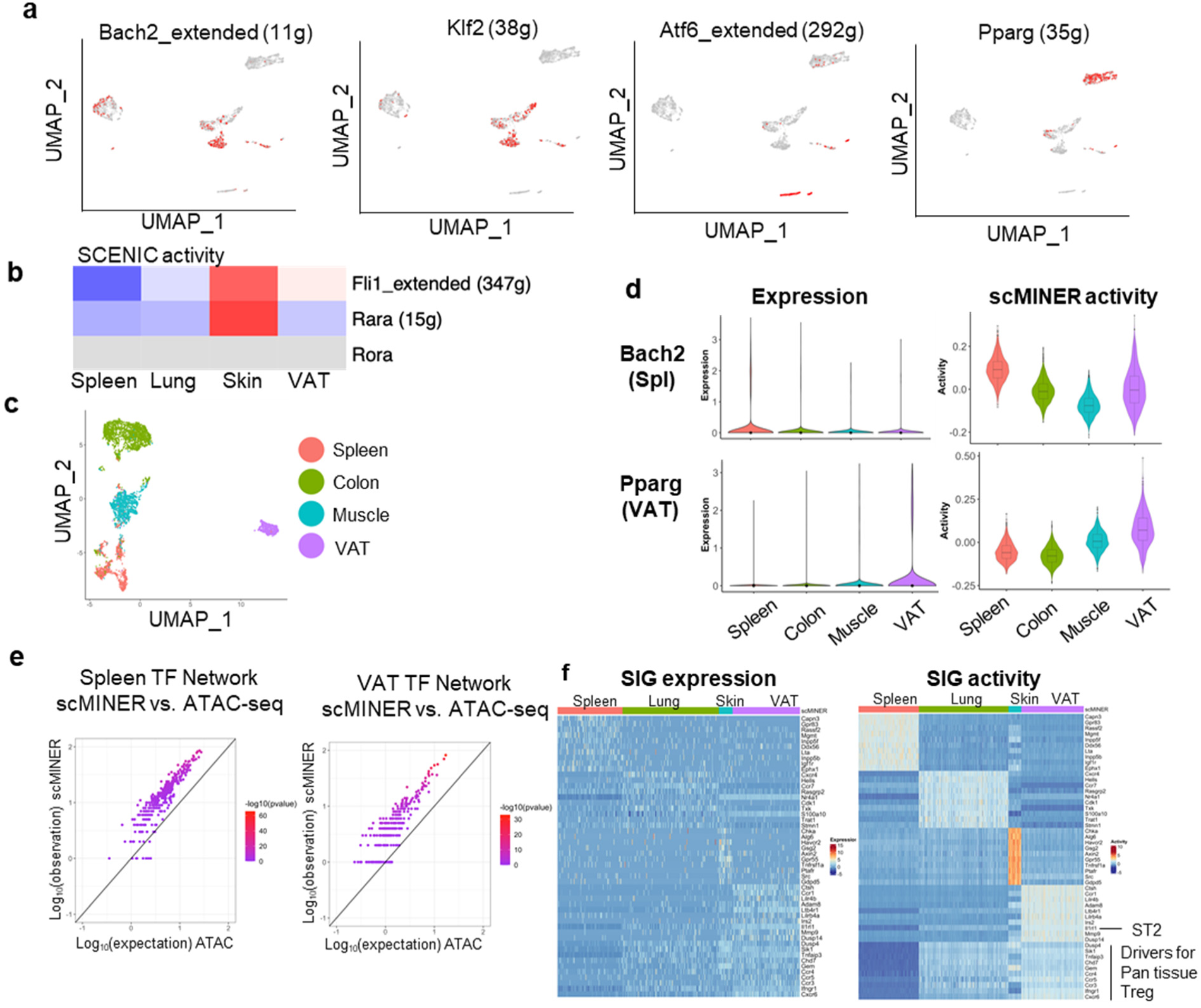
scMINER showed reproducibility in unravelling drivers in tissue specific Treg cells from different datasets. **a,** UMAP visualization of SCENIC binary activity of Bach2, Klf2, Atf6 and Pparg. **b,** Heatmap of average SCENIC activity of FLI1, RARA and RORA in Treg cells from each tissue. Grey indicates that the TF activity could not be predicated by SCENINC. **c,** MICA MDS clustering of mouse Foxp3^+^ regulatory CD4^+^ T cells (GSE109742) isolated from spleen, colon, muscle and visceral adipose tissue (VAT). **d,** Violin plot visualization of *Bach2* and *Pparg* expression and scMINER activity in spleen, colon, muscle and VAT Treg cells from GSE109742. **e,** Similarity of TF regulon in spleen and VAT Treg cells (GSE109742) generated by SJARACNe and footprint genes detected by ATAC-seq data (GSE112731) in corresponding tissues. Expected number of genes in intersection of ATAC-seq footprints as reference (log10 scale, x axis) with regard to hypergeometric distribution vs. observed intersection (log10 scale, y axis). For all genes, the observed intersection is significantly higher than expectation (black line). The color of the dots represents the -log_10_ (P-value) according to Fisher’s exact test. **f,** Heatmap visualization of SIG expression in each cell clustered by mouse Foxp3^+^ regulatory CD4^+^ T cells isolated from spleen, lung, skin and VAT. Drivers for Pan tissue Treg, drivers that have higher activity in Treg cells from the lung, skin and VAT than from spleen.

**Supplementary Table 1.** Summary of 11 single-cell datasets used for the evaluation of clustering methods.

| Dataset | Protocol | Size | Class | Taxonomy | Tissue | Accession ID |
| --- | --- | --- | --- | --- | --- | --- |
| Yan (2013)<sup>1</sup> | Tang | 124 | 8 | Human | Embryonic stem | GSE36552 |
| Goolam (2016)<sup>2</sup> | Smart-Seq2 | 124 | 5 | Mouse | Development | E-MTAB-3321 |
| Buettner | C1 | 182 | 3 | Mouse | Embryonic stem | E-MTAB-2805 |
| Pollen (2014)<sup>4</sup> | SMARTer | 301 | 11 | Human | Cerebral cortex | SRP041736 |
| Chung (2017)<sup>5</sup> | SMARTer | 515 | 5 | Human | Breast cancer | GSE75688 |
| Usoskin (2015)<sup>6</sup> | STRT-seq | 622 | 4 | Mouse | Sensory neurons | GSE59739 |
| Kolod (2015)<sup>7</sup> | SMARTer | 704 | 3 | Mouse | Embryonic stem | E-MTAB-2600 |
| Klein (2015)<sup>8</sup> | inDrop | 2,717 | 4 | Mouse | Embryonic Stem | GSE65525 |
| Zeisel (2015)<sup>9</sup> | STRT-seq | 3,005 | 7 | Mouse | Cortex, hippocampus | GSE60361 |
| Zheng (2017)<sup>10</sup> | 10x Genomics | 20,000 | 10 | Human | Sorted peripheral | SRP073767 |
| Bakken (2020)<sup>11</sup> | 10x Genomics | 76,533 | 20 | Human | Motor cortex | Azimuth |

**Supplementary Table 2.** Summary of scRNA-seq and ATAC-seq datasets used for scMINER applications.

| Accession ID | Data type | Cell types | Protocol |
| --- | --- | --- | --- |
| GSE122712 | scRNA-seq | CD8<sup>+</sup> T cells from chronic infection<sup>12</sup> | 10x Genomics |
| GSE130879 | scRNA-seq | Tissue (spleen, lung, skin, and VAT) Treg cells<sup>13</sup> | 10x Genomics |
| GSE130879 | scRNA-seq | Tissue Treg precursors<sup>13</sup> | 10x Genomics |
| GSE109742 | scRNA-seq | Tissue (spleen, colon, muscle, and VAT) Treg cells<sup>14</sup> | InDrop |
| GSE119940 | scRNA-seq | CD8<sup>+</sup> T cells from WT and Tox KO in chronic infection (day 7)<sup>15</sup> | 10x Genomics |
| GSE123236 | ATAC-seq | Tpex and Tex in LCMV infection<sup>12</sup> | Bulk |
| GSE132986 | ATAC-seq | WT and Tox KO CD8<sup>+</sup> T cells<sup>16</sup> | Bulk |
| GSE112731 | ATAC-seq | Tissue (spleen and VAT) Treg cells<sup>14</sup> | Bulk |
| GSE156112 | scATAC-seq | Tissue (spleen, lung, skin, and VAT) Treg cells<sup>17</sup> | 10x Genomics |

**Supplementary Note: Comprehensive scMINER documentation and tutorial with examples is publicly accessible via** https://jyyulab.github.io/scMINER.

